# Tracking the Evolutionary Patterns of RNA Modifications from Bryophyte to Flowering Plants by Mass Spectrometry

**DOI:** 10.1101/2025.08.15.670441

**Authors:** Zhongyan Zhou, Wenjun Xiao, Na Wang, Wen Su, Zhiyu Yang, Zhiru He, Siying Liu, Huafu Pei, Cheng Guo, Xinhong Guo, Lei Yue

**Author notes:** Correspondence: Cheng Guo,; Xinhong Guo,; Lei Yue. These authors contributed equally to this work (Zhongyan Zhou, Wenjun Xiao, Na Wang).

## Abstract

RNA modifications are critical for the regulation of gene expression. Comprehensive profiling of RNA modifications is a prerequisite for unravelling their identities and functions. The plant hormone auxin plays pivotal roles in regulating plant growth, development, and environmental responses. In contrast to the extensive studies in animals, there is a paucity of knowledge regarding the existence and functions of RNA modifications in plants. This is due to the low abundance of modified ribonucleosides and the lack of accurate analytical methods. This study employs a state-of-the-art hydrophilic interaction liquid chromatography-tandem-mass spectrometry (HILIC-MS/MS) platform to achieve the sensitive detection of 12 nucleosides at the pM to nM level. Utilizing this platform, we were able to identify *N*^4^,2’-O-dimethylcytidine (m^4^C_m_) in plants by LC-MS/MS for the first time, which provided direct evidence for the widespread of this modification in plants. Moreover, a comprehensive investigation was undertaken to examine the 12 types of RNA modifications and their response patterns to auxin in a range of prototype plants, spanning from bryophyte (*Marchantia polymorpha*) to flowering plants (*Arabidopsis thaliana, Oryza sativa*, and *Zea mays*). The evolutionary patterns of the 12 RNA modifications were depicted from bryophyte to flowering plants, and their potential functions in plant growth, development, and auxin responses were revealed.

## INTRODUCTION

RNA molecules are some of the most fundamental molecular components in all organisms. During transcription or post-transcription, the basic building blocks of DNA and RNA (adenine, uracil, guanine and cytosine) can be chemically modified by various enzymes, thereby expanding their chemical diversity^1-3^. The collection of RNA modifications, which introduce functional, regulatory, and structural complexity to gene expression regulation, is commonly referred to as the epitranscriptome^4^. In 2011, the *N*^6^-methyladenosine (m^6^A) demethylase FTO was identified, demonstrating the reversibility of m^6^A modification and initiating research into the epigenetic modification of RNA^5^. Moreover, the demethylation of m^6^A, mediated by FTO, has been demonstrated to regulate biomass and yield in rice and potato^6^. This has opened up new pathways for crop improvement and molecular breeding at the level of post-transcription modification. To date, over 170 modifications have been identified in RNA^3^. RNA modifications have been demonstrated to regulate a number of key physiological processes in plants, including signaling pathways, the cell cycle, flowering time, and stress responses, thereby affecting plant growth, development and environmental adaptability^6-11^. However, only a limited number of these have been documented in plants, with the majority of these studies focusing on *Arabidopsis thaliana* (*A. thaliana*) and *Oryza sativa* (rice)^8^.

Epitranscriptomic regulation of RNA plays a multifaceted role in plant early development and reproductive processes as well as stress responses^12-19^. Plant RNA modifications encompass a range of chemical modifications, including the methylation of bases [e.g., m^6^A, *N*^1^-methyladenosine (m^1^A), and 5-methylcytosine (m^5^C)], methylation of the ribose sugar [e.g., 2′-*O*-methylation (N_m_) and *N*^6^, 2′-*O*-dimethyladenosine (m^6^A_m_)], acetylation of bases [e.g., *N*4-acetylcytidine (ac^4^C)], and more complex reactions involving oxidation of methylated bases [e.g., 5-hydroxymethylcytosine (hm^5^C)]^20-23^. Nevertheless, the knowledge about RNA modifications in plant organogenesis and evolution remains limited.

The advent of highly specific, selective, and sensitive analytical methods, such as liquid chromatography-mass spectrometry (LC-MS), has led to the development of powerful approaches for identifying and characterizing RNA modifications^24-27^. Compared with alternative techniques such as thin-layer chromatography (TLC)^28^, capillary electrophoresis (CE)^29^, microarrays^30^, and next-generation sequencing^31^, LC-MS offers a direct detection of chemical modifications that alter the mass of the four canonical nucleosides, thereby providing direct evidence for the discovery of novel modifications^32-37^. The plant hormone auxin plays a crucial role in regulating numerous aspects of plant growth, development, and environmental responses^38^. The emerging evidence suggests that m^6^A modulates the response to auxin in *A. thaliana*, rice, and *Zea mays* (maize)^39-42^. However, there is still much to be discovered regarding to the responsive profiles of RNA modifications and their evolutionary patterns within the plant kingdom.

In this study, a high sensitivity mass spectrometry method, hydrophilic interaction liquid chromatography-tandem-mass spectrometry (HILIC-MS/MS)^43-45^ was employed to examine diverse RNA modifications in a range of plant species. These include bryophyte [*Marchantia polymorpha* (*M. polymorpha*)] and flowering plants (*A. thaliana*, rice and maize). Details of the modifications can be found in Table S1 and Fig.S1. Notable fluctuations in the 12 identified RNA modifications in response to 1-Naphthaleneacetic acid (NAA), a prototype plant hormone and analog of indole-3-acetic acid^46, 47^, have been observed. It is noteworthy that among the 12 identified RNA modifications, *N*^4^,2’-O-dimethylcytidine (m^4^C_m_), has been first identified by LC-MS/MS in a variety of plants. This finding extends the known range of this modification to include Eubacteria and mammals (Scheme 1, Table S2)^48^. In conclusion, we demonstrated the evolutionary patterns of the RNA modifications in plants, spanning from bryophyte to flowering plants, and their potential functions in plant growth, development, and auxin responses.

**Scheme 1.**
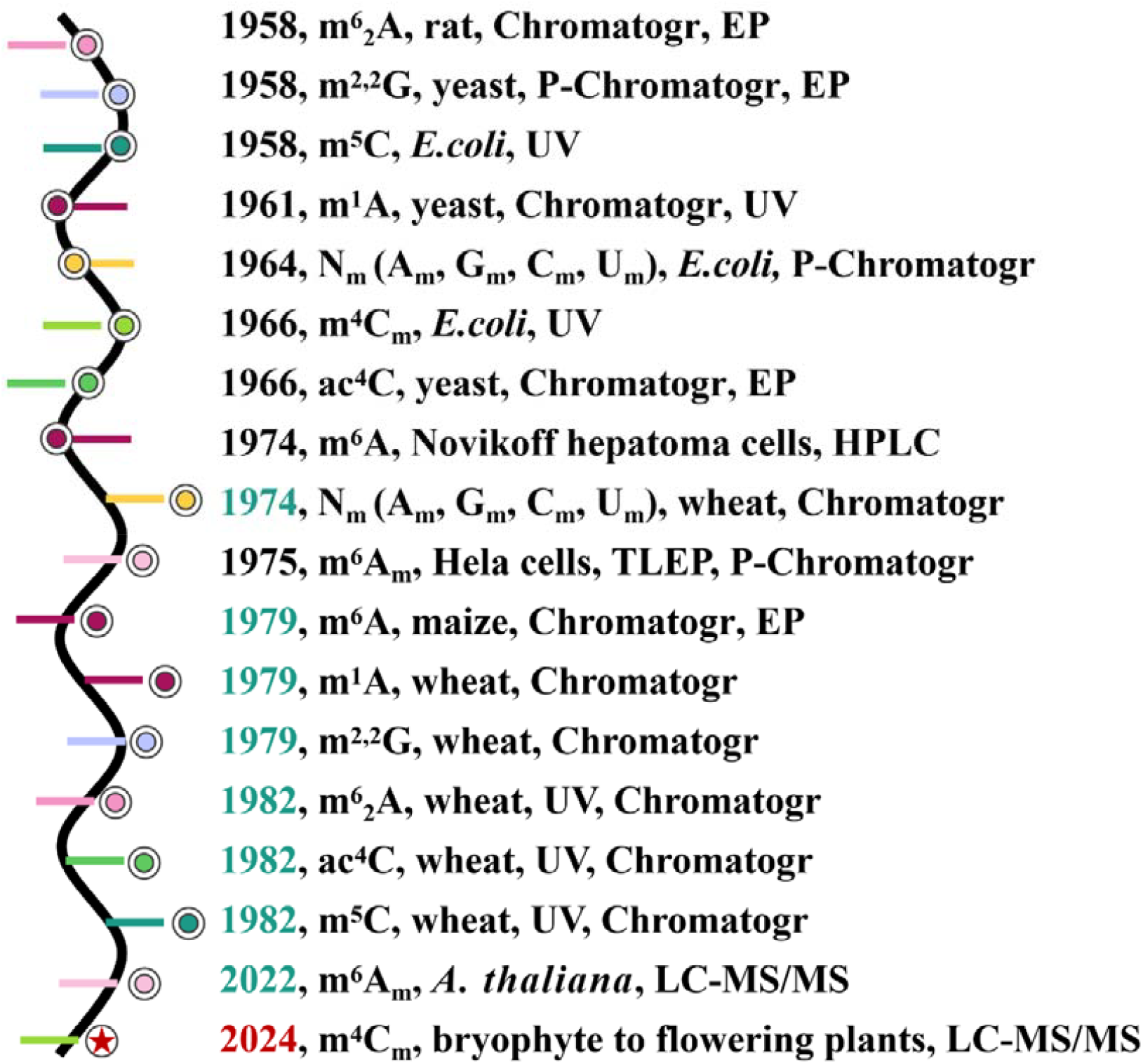
Timeline of identification and characterization of RNA modifications. The timeline illustrates the first reports of 12 RNA modifications detected in different species, highlighting the species and methods used for their identification. The data was sourced from Table S2 in the Supporting Information. The colors of the years represent different categories: black represents non-plant species, green represents previously identified modifications found in plants, and red represents RNA modification discovered in plants for the first time by LC-MS/MS in this study. Abbreviations: P-Chromatogr (paper chromatography), Chromatogr (chromatography), EP (electrophoresis), UV (ultraviolet spectral), HPLC (high performance liquid chromatography), TLEP (thin-layer electrophoresis).

## MATERIALS AND METHODS

### Chemical and Reagents

16 nucleoside standards and 14 stable isotope labeled internal standards were purchased from Sigma Aldrich or Toronto Research Chemical (Table S1). The purchasing information of other reagents and kits: ammonium formate, malic acid, agar, 1-naphthaleneacetic acid (NAA) and chloroform (Sigma Aldrich, USA), HPLC-grade acetonitrile (Merck, Germany), Formic acid (Fluka, Muskegon, MI), sucrose (Sinopharm, China), Nuclease P1 (NP1), erythro-9-(2-hydroxy-3-nonyl) adenine (EHNA), phosphodiesterase2 (PDE2), acetate, zinc chloride, antarctic phosphatasesodium (AP), antarctic phosphatase buffer (New England Biolabs, USA), 1/2 Murashige and Skoog medium (PM1061, Coolaber, China), Plant RNA Extraction Kit (19291ES50, Yeasen, China). Water was purified through the Milli-Q ultra-pure water system (Millipore, USA).

### Plant Materials and NAA Treatment

10-day seedlings of *A. thaliana* (Col-0) cultured on 1/2 Murashige and Skoog medium plates were treated with or without 100 ng/mL NAA for 30 min. The whole seedlings were collected for total RNA extraction. 10-day seedlings of rice (Nipponbare) cultured in Yoshida solution were treated with or without 100 ng/mL NAA for 30 min. The leaves and roots were collected respectively for total RNA extraction. 10-day seedlings of maize (B73) cultured in Yoshida solution were treated with or without 100 ng/mL NAA for 30 min. The leaves and roots were collected respectively for total RNA extraction. The plants of *M. polymorpha* (Tak1) cultured on 1/2 Murashige and Skoog plates were treated with or without 100 ng/mL NAA for 30 min. The thallus with rhizoids was collected for total RNA extraction.

### Isolation of Total RNA

The total RNA of *M. polymorpha, A. thaliana*, rice and maize was extracted and purified using the Plant RNA Extraction Kit according to the manufacturer’s recommended procedure. The concentrations were determined on a nanodrop 2000 spectrophotometer (Thermo Fisher Scientific, USA).

### Enzymatic Digestion of RNA

1 μg RNA was added 0.1 unit of nuclease P1, 0.25 nmol of EHNA, 0.000125 unit of PDE2, and a 3 μL solution containing 300 mM sodium acetate (pH 5.6) and 10 mM zinc chloride. Then water was added to the reaction mixture to reach a final volume of 15 μL. The reaction mixture was incubated at 37°C for 4 h. Then 0.5 units of AP, 3 μL 10× antarctic phosphatase buffer and water were added to reach a final volume of 30 μL. After digestion for 2 h at 37°C, the resulting digestion mixture was dried with Speed-vac and the dried residue was redissolved in 100 μL of water. Next, 25 μL aliquot of the digestion mixture of total RNA (250 ng) were spiked with 10 μL stable isotope labeled internal standards mixed solution (250 nmol [^13^C_5_]A, 1000 nmol [^13^C^15^N_2_]G), 500 nmol [^13^C_5_]C, 500 nmol [^13^C^15^N_2_]U, 0.625 nmol [D_3_]m^6^A_m_, 62.5 nmol [D_3_]U_m_, 123 nmol [D_3_]A_m_, 4 nmol [D_3_]m^6^A, 6.25 nmol [D_3_]m^1^A, 10 nmol [^13^C_5_]ac^4^C, 12.5 nmol [D_3_]C_m_, 6.25 nmol [D_6_]m^2,2^G, 62.5 nmol [D_3_]G_m_, 6.250 nmol [^13^CD_3_]m^5^C), 15 μL water and 50 μL chloroform, then the mixture was vortexed for 1 minute and centrifugation at 12000 rpm for 15 min to remove all enzymes used for the RNA digestion. The upper aqueous phase containing nucleosides was collected and dried. The dried residue was redissolved in 100 μL acetonitrile/water (9:1, v/v) to remove the salts. The solution was subsequently centrifuged, and 90 µL of the supernatant was dried. Subsequently, the dried sample was redissolved in 25 μL acetonitrile/water (9:1, v/v) for HILIC-MS/MS detection.

### HILIC-MS/MS Analysis

HILIC-MS/MS was performed on a 4000 QTRAP mass spectrometer (AB SCIEX, USA) equipped with an electrospray ionization source and a UPLC-MS/MS system consisting of an Acquity Primer UPLC system (Waters, USA). The samples were detected in the positive-ion mode. A Waters Primer BEH Amide column (2.1 mm × 100 mm, 1.7 μm) was employed for the chromatographic separation at 30°C. The samples were maintained at 4°C. A solution of 0.2% formic acid, 10 mM ammonium formate, 0.06 mM malic acid in water (mobile phase A) and 0.2% formic acid, 2 mM ammonium formate, 0.06 mM malic acid in acetonitrile (mobile phase B) were used as the mobile phases. The nucleosides were separated using a gradient of 95% B in 0-5.5 min, 95-92% B in 5.5-7 min, 92-85% in 7-9 min, 85-83% B in 9-9.5 min, 83-80% B in 9.5-11 min, 80% B in 11-13 min, 80-95% B in 13-13.5 min, and 95% B in 13.5-17 min. The flow rate was 0.3 mL/min. Each sample was injected 5 µL. In order to reduce interference, we used a switching valve to introduce the 0.5-14 min eluent from the column into the ion source.

Quantification of these nucleosides were performed in multiple reaction monitoring (MRM) mode. The ion transitions monitored for these modified nucleosides and the optimized MRM parameters were listed in the Table S7. The ion source temperature and spray voltage were set at 550°C and 5.5 kV, respectively. The curtain gas and ion source gases 1 and 2 were set at 40, 50 and 50 psi, respectively.

### Method Validation

The performance of the method was assessed on the basis of various parameters including selectivity, linearity, limit of detection (LOD), limit of quantification (LOQ), precision and accuracy.

Standard nucleosides solution of different concentration were mixed with fixed concentration of stable isotope labeled internal standards mixed solution (250 nmol [^13^C_5_]A, 1000 nmol [^13^C^15^N_2_]G), 500 nmol [^13^C_5_]C, 500 nmol [^13^C^15^N_2_]U, 0.625 nmol [D_3_]m^6^A_m_, 62.5 nmol [D_3_]U_m_, 123 nmol [D_3_]A_m_, 4 nmol [D_3_]m^6^A, 6.25 nmol [D_3_]m^1^A, 10 nmol [^13^C_5_]ac^4^C, 12.5 nmol [D_3_]C_m_, 6.25 nmol [D_6_]m^2,2^G, 62.5 nmol [D_3_]G_m_, 6.250 nmol [^13^CD_3_]m^5^C). The calibration curve was described as y = ax+b by measuring the ratio of the peak area of the analyte to the corresponding stable isotope labeled internal standard (y) with respect to the concentration of the analyte (x). For m^6^ A, [D_3_]m^6^A was used as the internal standard. Also, for m^4^C m, [^13^CD_3_]m^5^C was used as the internal standard. The LOD and LOQ were obtained by measuring the standard solutions at signal-to-noise ratios of 3 and 10, respectively. Intra- and inter-day precision was assessed by measuring three different levels of quality control (QC) samples in one day and on three consecutive days. The accuracy was assessed by comparing the measured concentrations of QC samples with their theoretical values.

### Statistical Analysis

The statistical analysis was carried out using the GraphPad Prism 8 software. The unpaired *t*-test was used to evaluate the RNA modifications in *M. polymorpha, A. thaliana*, rice, and maize between NAA treated and untreated plants. The *p*-value less than 0.05 was considered statistically significant.

## RESULTS AND DISCUSSION

### 1. Determination of Modifications in Total RNA by HILIC-MS/MS Analysis

The initial step was to establish a comprehensive HILIC-MS/MS method for the detection of RNA modifications. As shown in Fig.1, the separation was executed with remarkable efficacy under optimized chromatographic separation conditions. A total of 12 kinds of modifications (m^6^A_m_, U_m_, m^6^ A, A_m_, m^6^A, m^1^A, m^4^C_m_, ac^4^C, C_m_, m^2,2^G, G_m_ and m^5^C) and 4 kinds of nucleosides (A, U, C, and G) were identified and characterized. The characteristic extracted-ion chromatograms and the identical retention times from the total RNA of disparate samples align with those of the standard nucleosides (Figs.S5-S10), thereby substantiating the accuracy of the examination of the 12 modifications. Note that effective separation of m^1^A and m^6^A, a nucleosides isomer pair with exactly the same precursor ion and daughter ion in MRM detection, is achieved. The established method was then employed to detected modifications in the total RNA from *M. polymorpha, A. thaliana*, rice, and maize.

**Fig. 1.**
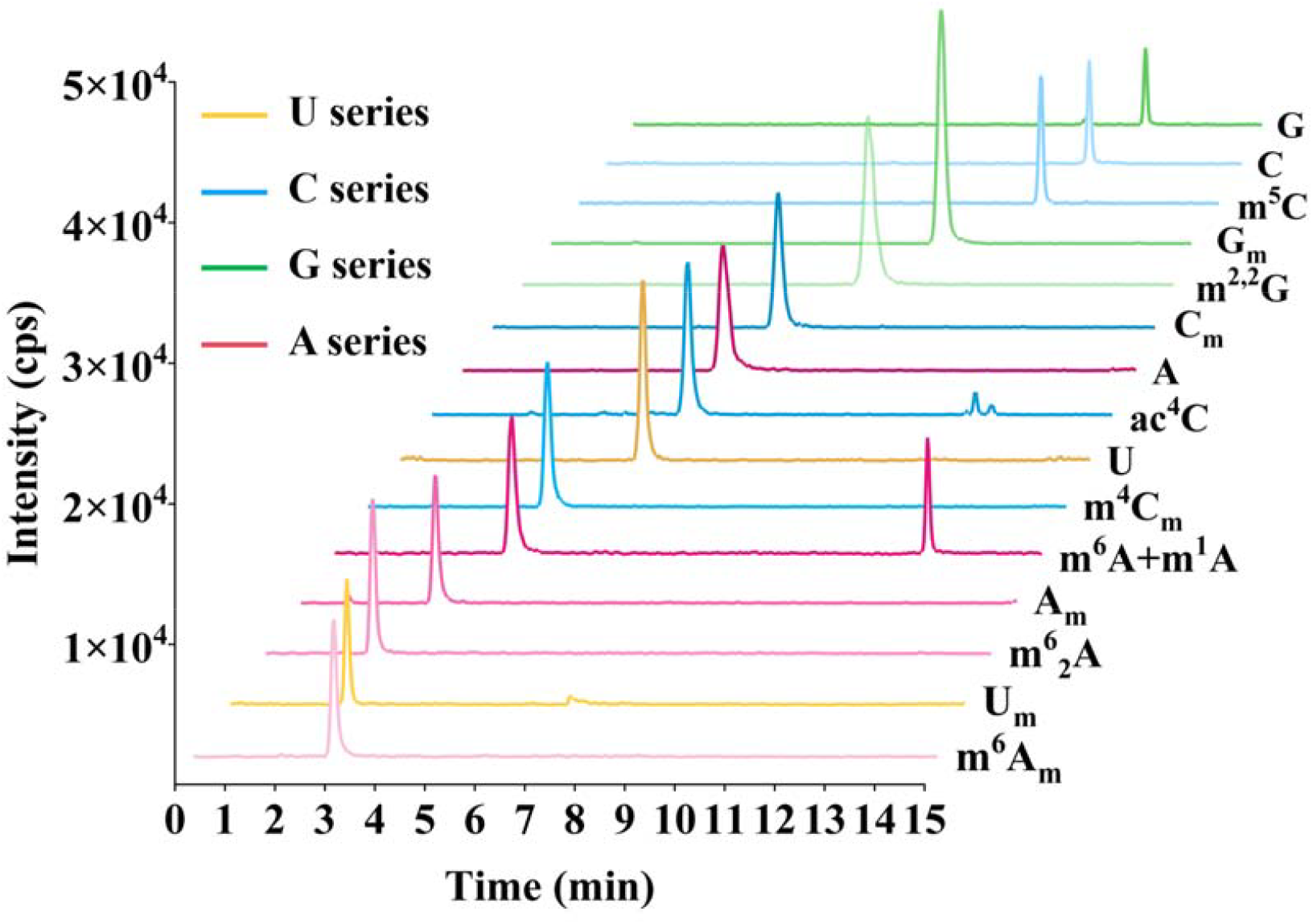
The MRM chromatograms of rA, rC, rG, rU, m^6^A_m_, U_m_, m^6^ A, A_m_, m^6^A,m^1^A, m^4^C_m_, ac^4^C, C_m_, m^2,2^G, G_m_, and m^5^C. Concentration: rU (250 nM), rA and U_m_ (50 nM), rG (25 nM), G_m_ (12.5 nM), m^2,2^G, m^5^C and ac^4^C (2.5 nM), A_m_, and C_m_ (1.25 nM), m^6^A and m^1^A (1 nM), m^6^_2_A and m^6^A_m_ (0.5 nM).

The application of the HILIC-MS/MS platform facilitated the direct identification of a previously uncharacterized RNA modification by LC-MS/MS, m^4^C_m_, in plants. In *M. polymorpha, A. thaliana*, rice and maize, a peak with a retention time of 3.9 minutes in the extracted-ion chromatogram (m/z 272.1→126.0), corresponding to the m^4^C_m_ standard (Fig.2), confirming the presence of m^4^C_m_. On the contrary, m^4^C_m_ was not detectable in the control sample of water or the sample with only adding enzymes (Fig.2), which excludes the possibility that the detected m^4^C_m_ was from the contamination of water or enzymes. This finding suggests that the detected m^4^C_m_ modification was from bryophyte and flowering plants. We have found that the m^4^C_m_ content in rice and maize leaves was higher than that in roots, indicating a potential role in leaf growth and development. Notably, the m^4^C_m_ signal in maize roots was merely the baseline and was thus identified as not detected, indicating that m^4^C_m_ modification may not be involved in maize root development. The aforementioned profiles indicate plausible spatial patterns and functional differentiation of m^4^C_m_ during organogenesis and plant evolution.

**Fig. 2.**
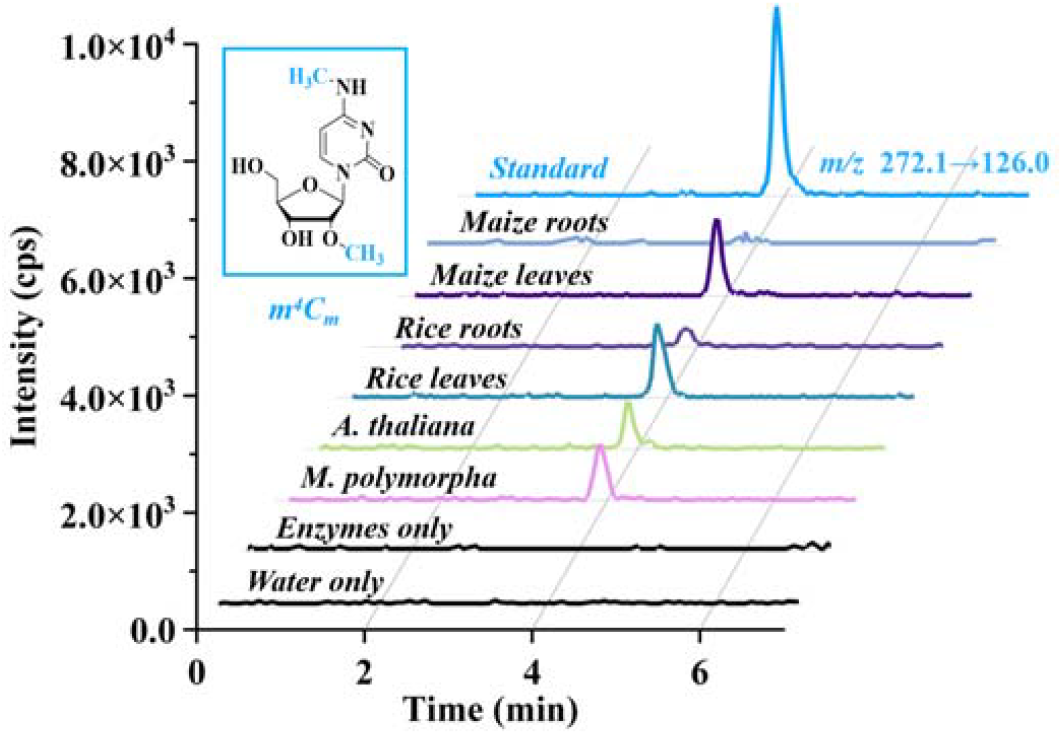
Determination of m^4^C_m_ in plants total RNA. The extracted-ion chromatograms of m^4^C_m_ in *M. polymorpha, A. thaliana*, rice leaves, rice roots, maize leaves, and maize roots, compared with the nucleoside standard. The chemical structure of m^4^C_m_. is shown in the upper left. Water or enzymes control samples represent the samples only containing water or enzymes used for digesting RNA.

RNA modifications play key roles in regulating gene expression, maintaining RNA stability and participating in plant growth and development^49, 50^. To further dissect the distribution of RNA modifications in different plants and their potential biological functions, the calibration curves were established for the 12 kinds of modifications (m^6^A_m_, U_m_, m^6^_2_A, A_m_, m^6^A, m^1^A, m^4^C_m_, ac^4^C, C_m_, m^2,2^G, G_m_, and m^5^C). The calibration curves demonstrated good linearity, with the coefficients of determination (R^2^) exceeding 0.99 (Table S3). Subsequently, a comprehensive quantification of RNA modifications was conducted in model plants, spanning from bryophyte (*M. polymorpha*) to dicotyledons (*A. thaliana*), and monocotyledons (rice and maize) (Fig.3 and S2).

**Fig. 3.**
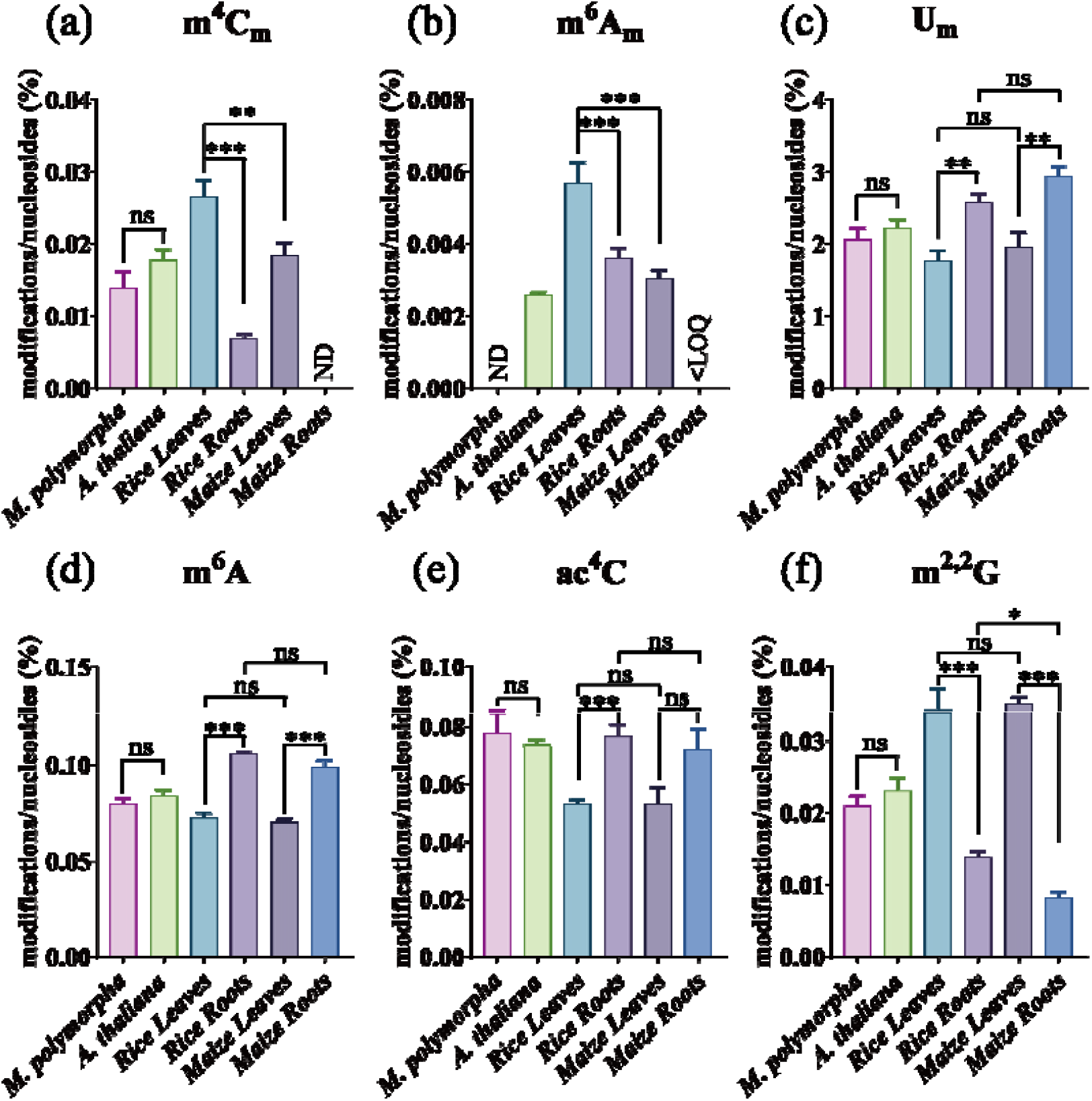
The quantification of (a) m^4^C_m_, (b) m^6^A_m_, (c) U_m_, (d) m^6^A, (e) ac^4^C, (f) m^2,2^G, in *M. polymorpha, A. thaliana*, rice leaves, rice roots, maize leaves and maize roots. <LOQ, below the instrumental limit of detection. ND, not detected. Data presents the mean ± standard error of the mean (SEM) of three independent experiments. Statistical analysis was performed by using unpaired *t*-test. * *p* < 0.05, ** *p* < 0.01; *** *p* < 0. 001, no significance.

Our findings indicate that m^6^A_m_ was detected in *A. thaliana*, rice, and maize, but the content of m^6^A_m_ in *M. polymorpha* was lower than the LOD (Fig.3b). And the abundance of m^1^A in *M. polymorpha* was observed to be lower than that in *A. thaliana* (Fig.S2c, suggesting that m^6^A_m_ and m^1^A emerged during the plant terrestrial evolution and may play a role in regulating the growth, development, and environmental adaptation of flowering plants, as opposed to bryophyte or lower plants. Conversely, the abundance of A_m_ in *M. polymorpha* was higher than that in *A. thaliana* (Fig.S2a), implying that A_m_ may play a more pivotal role in the adaptation of bryophytes to environments. Furthermore, the levels of nine additional RNA modifications (U_m_, m^6^_2_A, m^6^A, m^4^C_m_, ac^4^C, C_m_, m^2,2^G, G_m_, and m^5^C) were found to be comparable between *A. thaliana* and *M. polymorpha*. This observation suggests that these modifications may play an evolutionarily conserved and fundamental role in diverse biological processes within the plant kingdom.

Additionally, our findings indicate that the profiles of RNA modifications vary across different plant species. The results showed that the abundance of m^4^C_m_, m^6^A_m_ and m^5^C in rice leaves were higher than that in maize leaves (Figs.3a-b and S2f). Conversely, rice roots contained higher accumulation of m^2,2^G, m^1^A and m^5^C than maize roots (Figs.3f, S2c, and S2f). This suggests that rice may be more reliant on m^4^C_m_, m^6^A_m_, m^5^C, m^2,2^G and m^1^A for the regulation of leaves and roots development, in comparison to maize. In rice, ac^4^C was observed to accumulate preferentially in roots other than leaves, suggesting that ac^4^C plays a more pivotal role in rice roots development (Fig.3e). Other RNA modifications exhibited no appreciable differences in tissue distribution between rice and maize, indicating that these modifications have undergone conserved evolutionary patterns in monocots.

It is noteworthy that certain RNA modifications demonstrated a distinct tissue-specificity. The content of m^4^C_m_, m^6^A_m_, and m^2,2^G in both rice and maize leaves were significantly higher than that in roots (Figs.3a-b and 3f), while the abundance of U_m_, m^6^A, A_m_, m^6^ A, m^1^A, C_m_, and G_m_ in roots were remarkably higher than that in leaves (Figs.3c-d, 3f, S2a-c, and S2d-e). This suggests that different RNA modifications play diverse roles in the development of leaves and roots.

### 2. The Response of RNA Modifications to Auxin

To elucidate the relationship between RNA modifications in responses to environments, we employed NAA, a prototype auxin, to act as a possible environmental stimulation factor^38^. This allowed us to investigate how RNA modifications responded to NAA in plants. Following NAA treatment, notable alterations in RNA modification levels were discerned across all plants (Fig.4, Fig.S3-S4 and Tables S5-S6). With regard to the m^4^C_m_ modification, *A. thaliana* exhibited an inhibited level in response to NAA treatment, whereas rice and maize displayed opposite responsive patterns (Fig.4a). This suggests that the functional association between m^4^C_m_ and auxin signaling diverged among monocots and dicots. Interestingly, *M. polymorpha* exhibited a reduction in the level of G_m_ modification, whereas *A. thaliana*, rice, and maize demonstrated an increase in accumulation (Fig.4b), indicating that auxin treatment inhibited G_m_ modification of total RNA in bryophytes, but promoted it in flowering plants. Furthermore, m^6^A_m_ was not detected only in *M. polymorpha*, irrespective of whether the sample was taken before or after NAA treatment (Fig.4c), suggesting that angiosperms evolved more complex RNA modifications than bryophytes. As to ac^4^C, m^6^_2_A, m^1^A, C_m_, and m^5^C (Figs.4d-e, Fig.S3a-c), auxin was observed to exert a negative regulatory effect on the level of these five modifications in rice, while stimulating their accumulation in maize. These findings elucidate the multifaceted impact of auxin on RNA modifications in monocotyledonous plants. It is noteworthy that maize exhibited pronounced alterations in m^2,2^G, m^1^A, A_m_, and m^6^A compared to other plants, and displayed disparate U_m_ responses in its leaves and roots (Fig.4f, Fig.S3a and Figs.S3d-f), suggesting their pivotal involvement in maize growth, development, and auxin response.

**Fig. 4.**
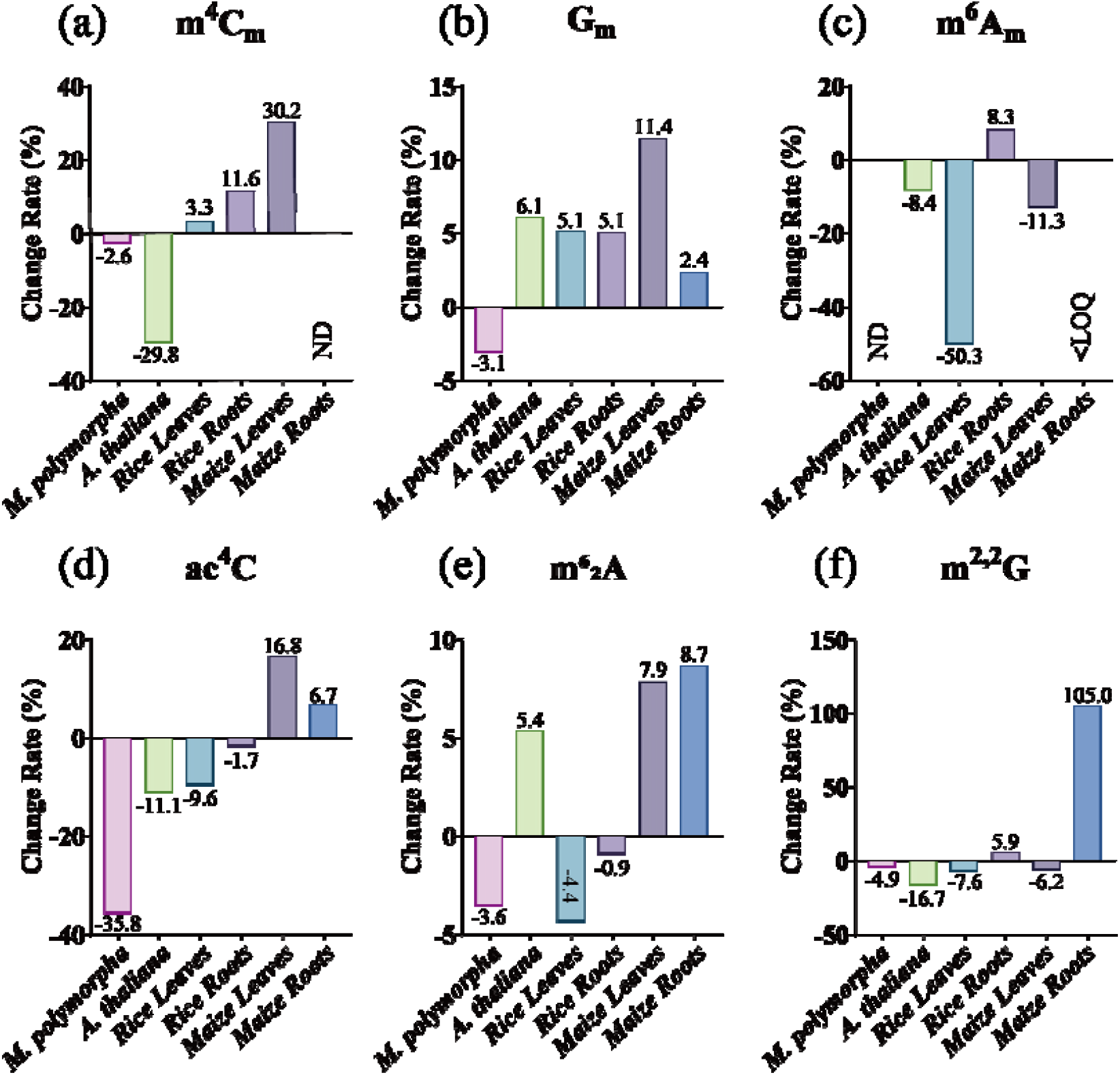
The change rate of (a) m^4^C_m_, (b) G_m_, (c) m^6^A_m_, (d) ac^4^C, (e) m^6^_2_A, (f) m^2,2^G, after NAA treatment in *M. polymorpha, A. thaliana*, rice leaves, rice roots, maize leaves and maize roots compared to the control. <LOQ, below the instrumental limit of detection. ND, not detected. The percentage of change rate was calculated using the following formula: change rate = (level of X after NAA treatment - level of X in the control) / level of X in the control × 100%, the X represents a specific modification.

In order to dissect the global response profiles to auxin, we conducted a further comparison of the 12 modifications that underwent change with and without NAA treatment (Fig.5). In *M. polymorpha*, U_m_ and m^1^A were significantly induced by auxin, while the other nine modifications were inhibited, particularly ac^4^C, C_m_ and m^5^C (Fig.5a). This indicates that the U_m_ modification may serve as a crucial inhibitor to the post-transcriptional or translational regulation of auxin responses, whereas the ac^4^C, C_m_ and m^5^C appear to exert opposing effects. In *A. thaliana*, five modifications (U_m_, G_m_, m^6^ 2A, m^6^A, and A m) are positively regulated, while seven modifications (m^4^C m, m^5^C, m^2,2^G, ac^4^C, m^6^_2_A, m^6^A_m_, C_m_, and m^1^A), are negatively regulated by auxin, respectively (Fig.5b). Interestingly, U_m_ is also the core regulator with the most significantly positive response to auxin, as observed in *M. polymorpha*. Furthermore, the response of the 12 modifications in rice leaves and roots exhibited distinct patterns (Figs.5c-d). The levels of G_m_ and m^4^C_m_ were elevated in both roots and leaves samples. The other 10 modifications decreased in the leaves, whereas m^6^A_m_, U_m_, m^2,2^G, G_m_ and A_m_ demonstrated an increase in the roots. In contrast to the findings in rice, the majority of RNA modifications in maize were up-regulated by auxin (Figs. 5e-f). It is noteworthy that m^2,2^G and U_m_ showed completely opposite responsive patterns in maize leaves and roots. On the whole, these results reveal the versatile roles of different RNA modifications in growth and development, as well as their spatial response patterns to auxin.

**Fig. 5.**
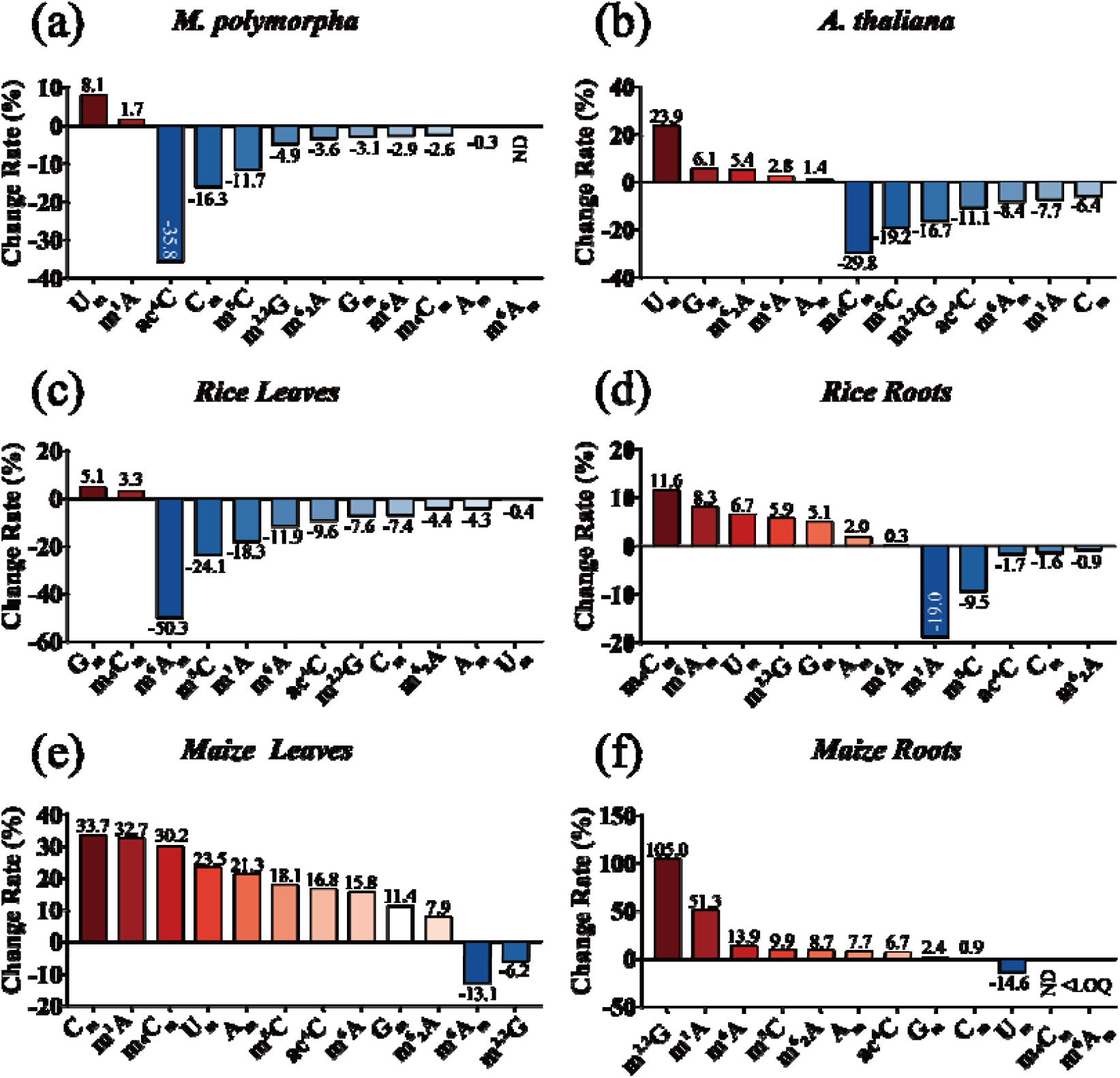
The change rate of 12 modifications in (a) *M. polymorpha*, (b) *A. thaliana*, (c) rice leaves, (d) rice roots, (e) maize leaves and (f) maize roots after NAA treatment compared to the control. <LOQ, below the instrumental limit of detection. ND, not detected. The percentage of change rate was calculated using the following formula: change rate = (level of X after NAA treatment - level of X in the control) / level of X in the control × 100%, the X represents a specific modification.

In this study, an ultra-high sensitivity HILIC-MS/MS analysis platform was employed to characterize the modification profiles of total RNA in *Marchantia polymorpha, Arabidopsis thaliana, Oryza sativa*, and *Zea mays*. 11 modifications were detected in *M. polymorpha*, and 12 modifications in *A. thaliana*, rice, and maize, respectively.

This finding indicates that *M. polymorpha* evolved a relatively comprehensive RNA modification system. It is noteworthy that m^4^C_m_, was identified and found to be widely present in bryophytes and flowering plants. Furthermore, the prevalence of RNA modifications exhibited notable variation across different plants and tissues. The observed diversities not only provide insights into the patterns of species evolution, but also reveal that RNA modifications could be closely related to the unique physiological and developmental needs of each species.

In further tissue-level analyses, RNA modifications in the roots and leaves of rice and maize showed different accumulation profiles, indicating the divergence of post-transcriptional modifications and functions in monocots. Furthermore, a systematic investigation was conducted to ascertain the response patterns of 12 RNA modifications to auxin. The three flowering plants *A. thaliana*, rice, and maize exhibited diverse responsive profiles, whereas *M. polymorpha*, a model plant in the transition from aquatic to terrestrial environments, displayed a weaker response to auxin. This evidence demonstrates that the crosstalk between RNA modifications and auxin signaling is emerging in *M. polymorpha*. The construction of the crosstalk also appears to be vital for the flowering plants. The future studies could focus on lower plants such as charophyte algae and chlorophyte algae, which are likely to contain more original information about the early evolution of RNA modifications, and dissect the crosstalk with plant hormone abscisic acid (ABA) that emerges during the aquatic to terrestrial transition to gain tolerance for abiotic stresses, such as drought, radiation, and extreme temperature.

A number of modifications have been identified as playing a crucial role in the auxin response, including U_m_ and ac^4^C in *M. polymorpha*, U_m_ and m^4^C_m_ in *A. thaliana*, G_m_, m^4^C_m_, and m^6^A_m_ in rice, C_m_, m^4^C_m_, and m^2,2^G in maize. Note that it has long been assumed that modifications in rRNA are mostly added in the initial rRNA processing and maturation steps and seem not to be a dynamic moiety within the host rRNA molecule due to their high conservation and low turnover of rRNA^20^. However, this assumption is being challenged by their response profiles to auxin. Notably, among the four model plants, the levels of A_m_, G_m_, U_m_, and C_m_ were considerably higher than that of m^6^A, the most extensively studied RNA modification, and the four exhibited a pronounced response to auxin. To this assumption, we will seek to elucidate the global functions of rRNA modifications in plant growth and development.

Finally, this study immediately highlights the necessity for the identification of the enzymes responsible for writing, reading, and erasing the newly discovered modifications. Enhancing our comprehension of the roles of these modifications in plant growth and development hinges on the development of methods with high precision and resolution for mapping the precise modification positions in target genes. However, the extremely low abundance and dynamic changes of RNA modifications present a formidable challenge to this endeavor.

## Credit Authorship Contribution Statement

**Zhongyan Zhou:** Writing original draft, Validation, Software, Project administration, Methodology, Investigation, Formal analysis, Data curation.

**Wenjun Xiao:** Writing original draft, Project administration, Methodology, Investigation, Formal analysis, Data curation.

**Na Wang:** Writing original draft, Project administration.

**Wen Su, Zhiyu Yang, Zhiru He, Siying Liu, and Huafu Pei:** Methodology, Investigation and Software.

**Cheng Guo:** Writing, review & editing, Supervision, Funding acquisition. **Xinhong Guo:** Writing, review & editing, Supervision, Funding acquisition. **Lei Yue:** Writing, review & editing, Supervision, Funding acquisition.

## Acknowledgments

We thank Prof. Tao Yang (Sichuan Agricultural University) and Prof. Duoyan Rong (Hunan University of Technology) for kindly providing *Marchantia polymorpha* (tak1) and *Zea mays* (B73) plants. This work is supported by grants from the National Natural Science Foundation of China (22174037, 22176167, 32300456, 32372124 and 32372128), and National Key Research and Development Program of China (2023YFF0600040).

## Supporting Information description

Information of nucleosides and their isotopic labeled standards; calibration curves, LODs and quantification LOQs; accuracy and precision; measured average levels of RNA modifications; the MRM transitions and optimal parameters of mass spectrometry; the quantification of RNA modifications; and identification of nucleosides in plants.

