## Supporting Information for "Tracking the Evolutionary Patterns of RNA Modifications from Bryophyte to Flowering Plants by Mass Spectrometry"

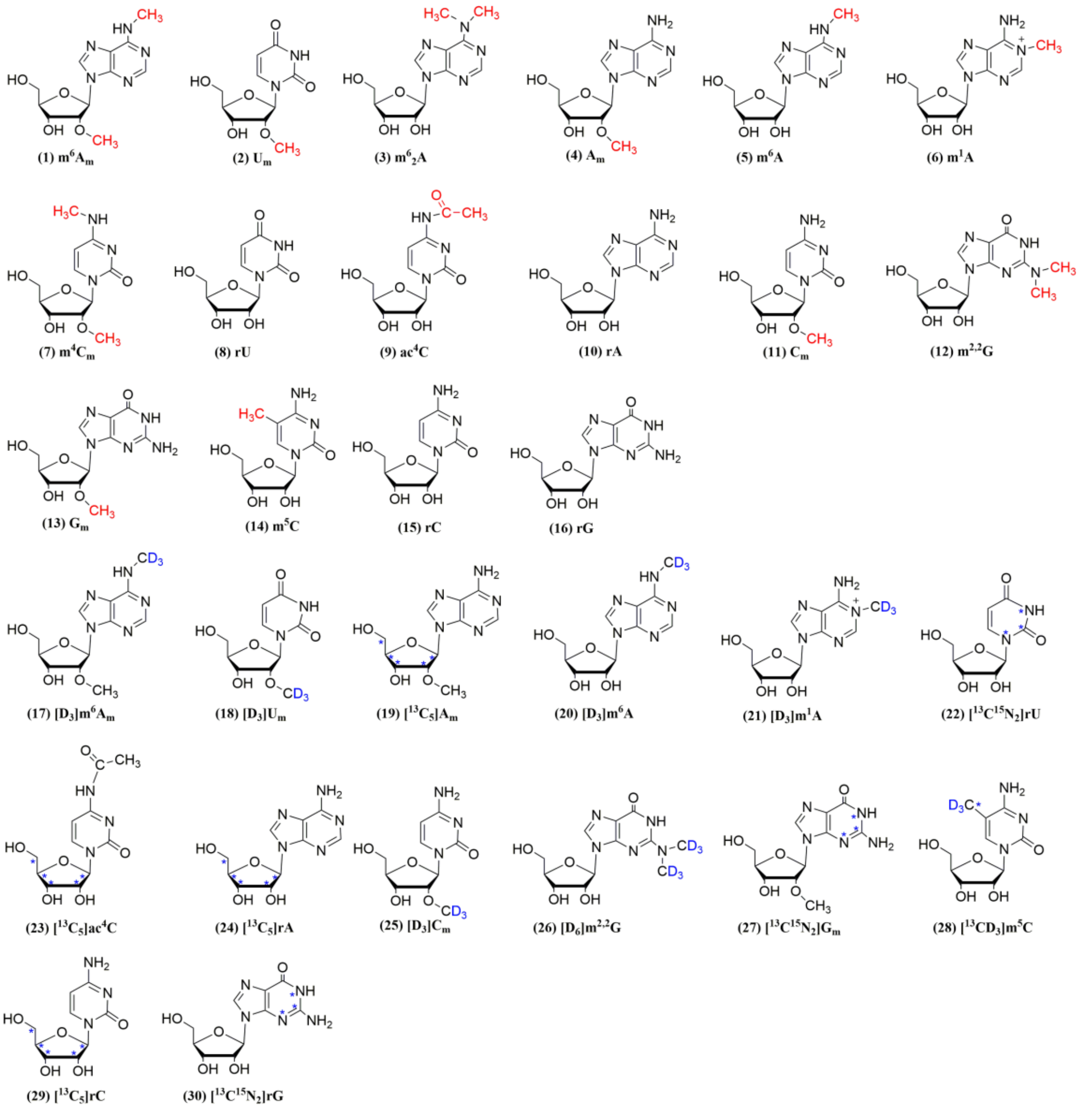


**Fig. S1** The chemical structures of $m^6A_m$, $U_m$, $m^6{}_2A$, $A_m$, $m^6A$, $m^1A$, $m^4C_m$, rU, $ac^4C$, rA, $C_m$, $m^{2,2}G$, $G_m$, $m^5C$, rC, and rG (red), and their stable isotope-labeled internal standards (blue).

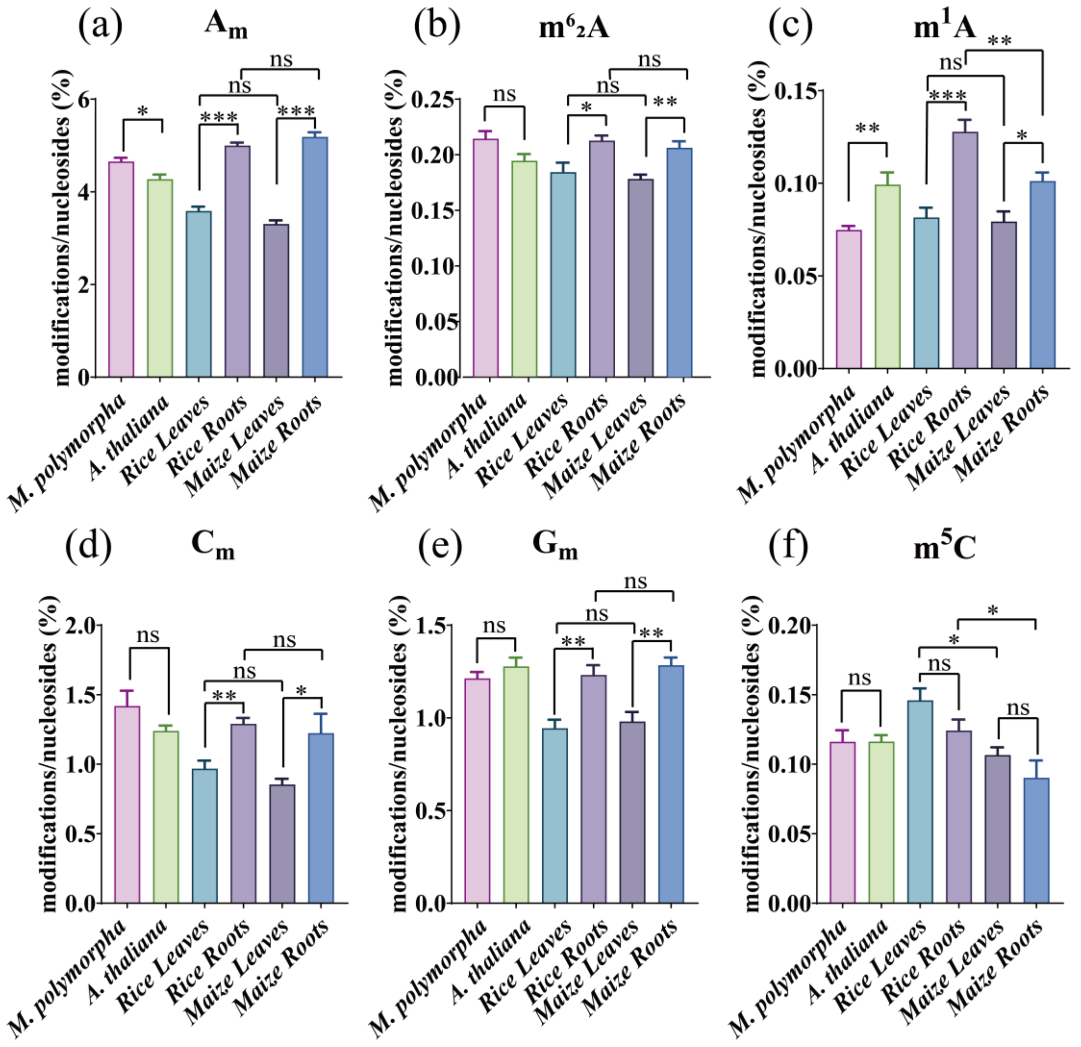


**Fig.S2** The quantification of (a) $A_m$, (b) $m^6{}_2A$, (c) $m^1A$, (d) $C_m$, (e) $G_m$ and (f) $m^5C$ in *M. polymorpha*, *A. thaliana,* rice roots, rice leaves, maize roots, and maize leaves. Data present the mean ± standard error of the mean (SEM) of three independent experiments. Statistical analysis was performed by using unpaired *t*-test. $p > 0.05$, * $p < 0.05$, ** $p < 0.01$, *** $p < 0.001$, ns, no significance.

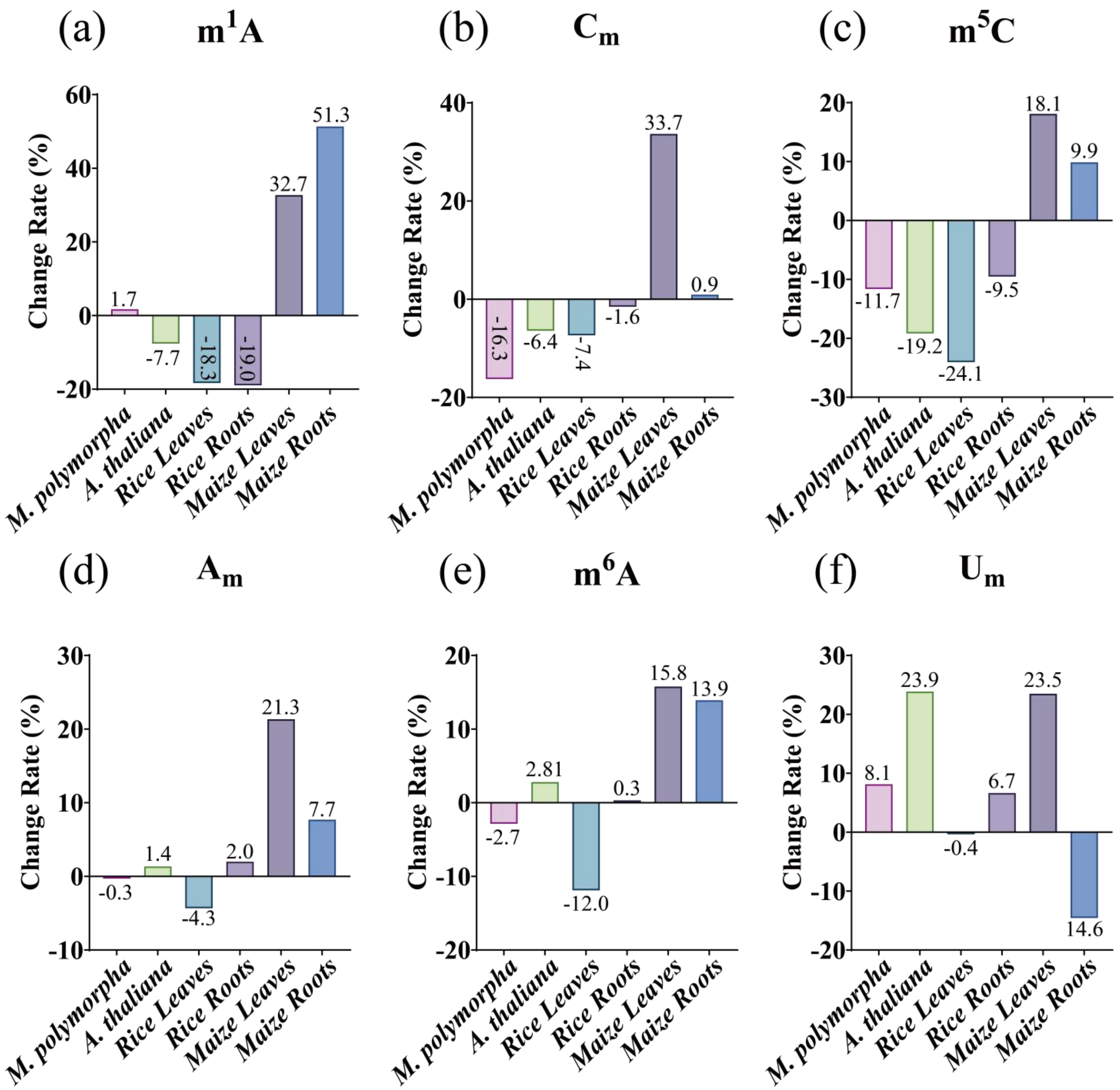


**Fig.S3** The change rate of (a) $m^1A$, (b) $C_m$, (c) $m^5C$, (d) $A_m$, (e) $m^6A$ and (f) $U_m$ in total RNA after NAA treatment in *M. polymorpha*, *A. thaliana*, rice leaves, rice roots, maize leaves and maize roots compared to the control. The percentage of change rate was calculated using the following formula: change rate = (level of X after NAA treatment - level of X in the control) / level of X in the control × 100%, the X represents a specific modification.

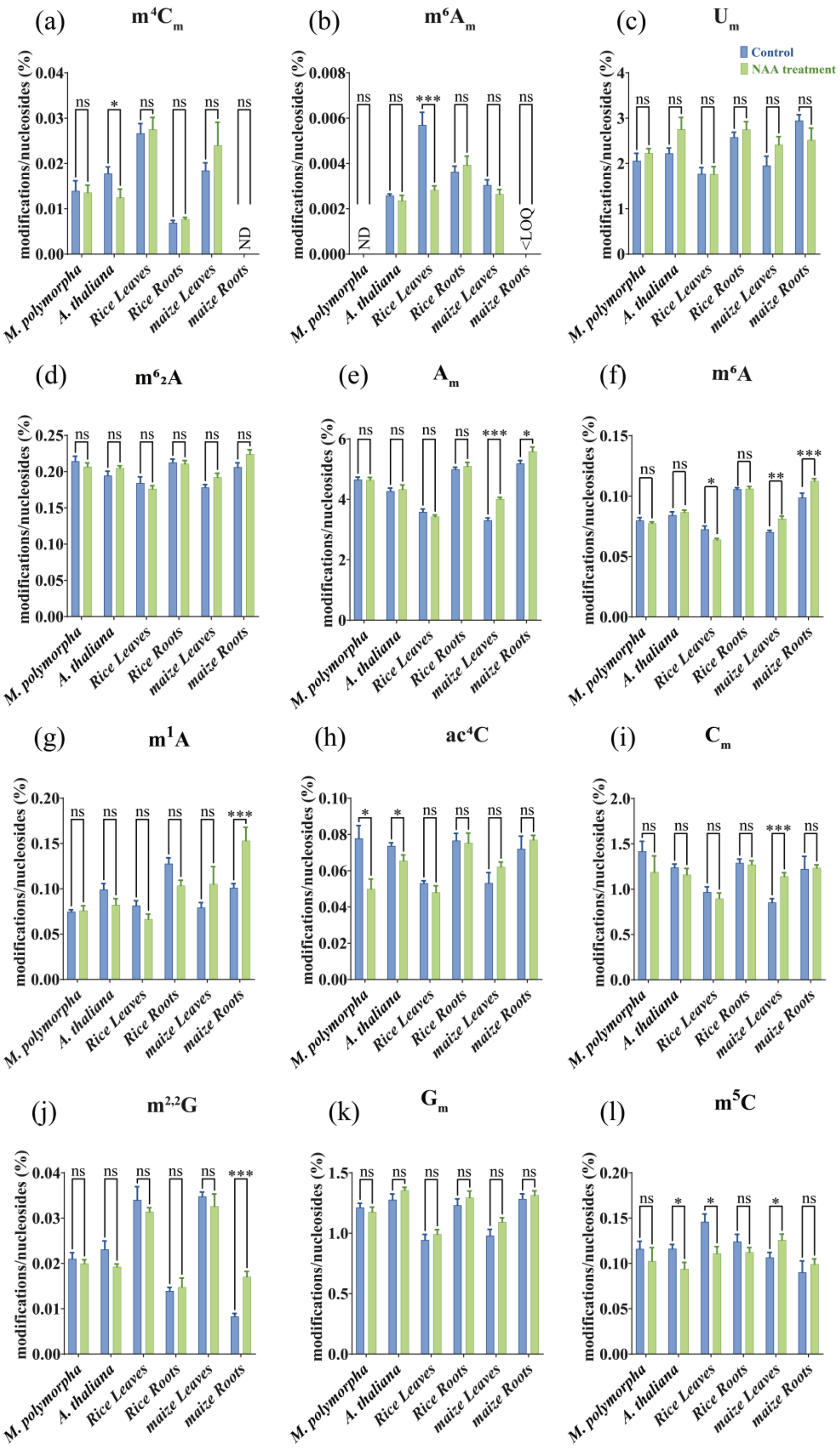


**Fig.S4** Quantification of the change of (a) $m^4C_m$, (b) $m^6A_m$, (c) $U_m$, (d) $m^6{}_2A$, (e) $A_m$, (f) $m^6A$, (g) $m^1A$, (h) $ac^4C$, (i) $C_m$, (j) $m^{2,2}G$, (k) $G_m$ and (l) $m^5C$ in total RNA upon NAA treatment. Data present the mean ± standard error of the mean (SEM) of three independent experiments. Statistical analysis was performed by using unpaired *t*-test. $p > 0.05$, * $p < 0.05$, ** $p < 0.01$, *** $p < 0.001$, ns, no significance.

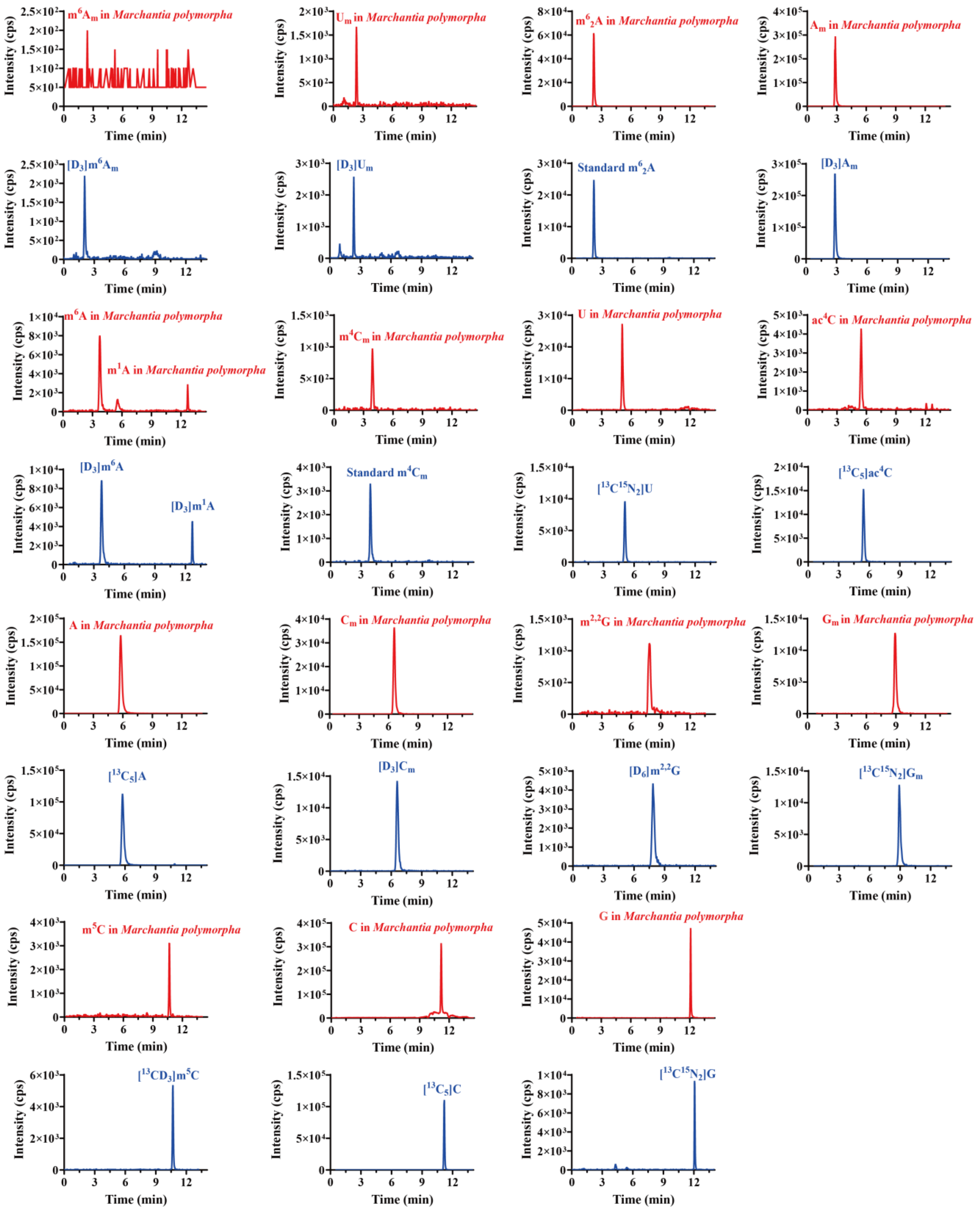


**Fig.S5** Identification of 16 kinds of nucleosides ($m^6A_m$, $U_m$, $m^6_2A$, $A_m$, $m^6A$, $m^1A$, $m^4C_m$, rU, $ac^4C$, rA, $C_m$, $m^{2,2}G$, $G_m$, $m^5C$, rC, and rG) in *Marchantia polymorpha* samples by HILIC-MS/MS analysis. The extracted-ion chromatograms of the modifications detected in total RNA of *Marchantia polymorpha* samples are highlighted in red and the spiked stable isotope-labeled internal standards are highlighted in blue.

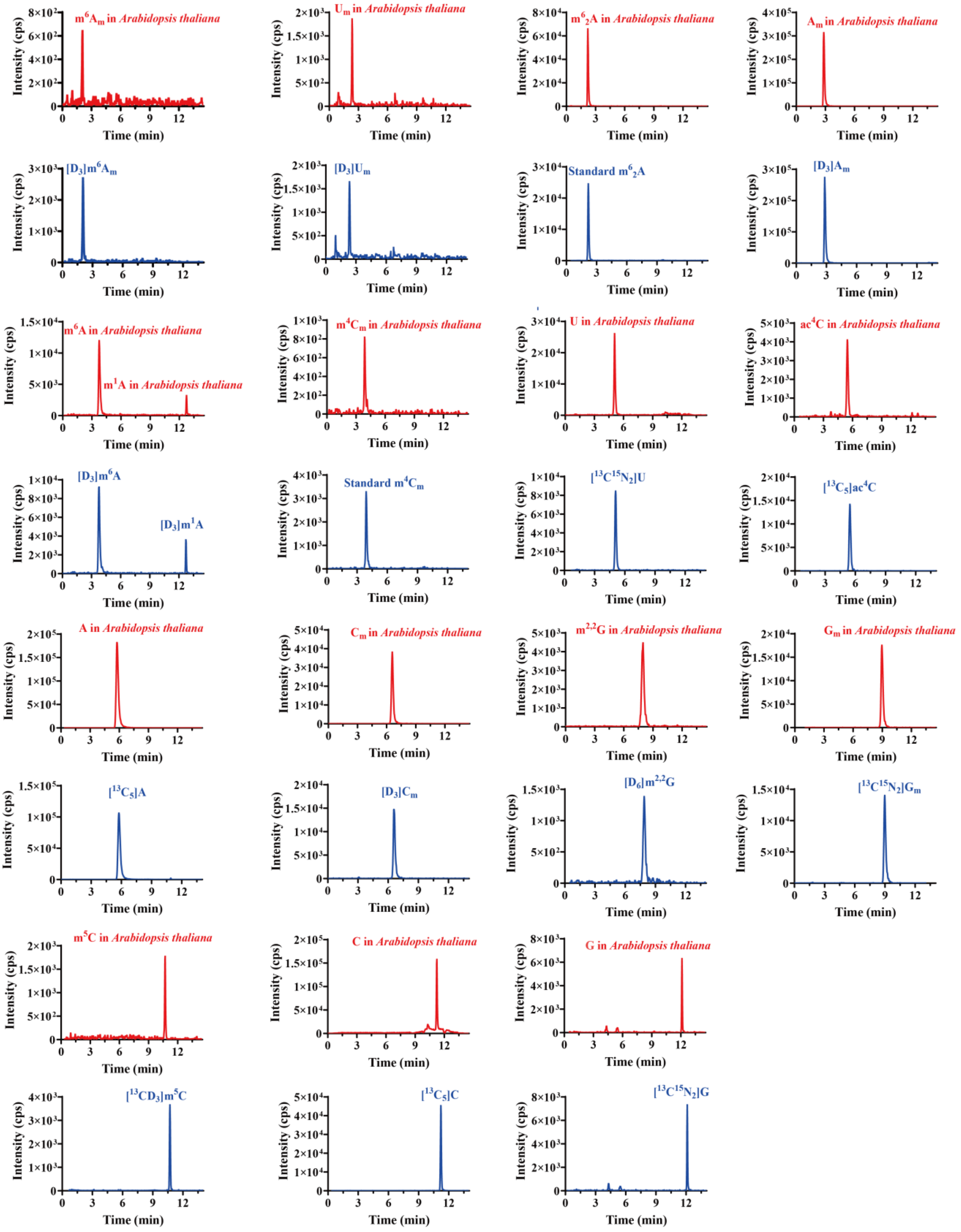


**Fig. S6** Identification of 16 kinds of nucleosides ($m^6A_m$, $U_m$, $m^6{}_2A$, $A_m$, $m^6A$, $m^1A$, $m^4C_m$, rU, $ac^4C$, rA,$C_m$, $m^{2,2}G$, $G_m$, $m^5C$, rC, and rG) in *Arabidopsis thaliana* samples by HILIC-MS/MS analysis. The extracted-ion chromatograms of the modifications detected in total RNA of *Arabidopsis thaliana* samples are highlighted in red and the spiked stable isotope-labeled internal standards are highlighted in blue.

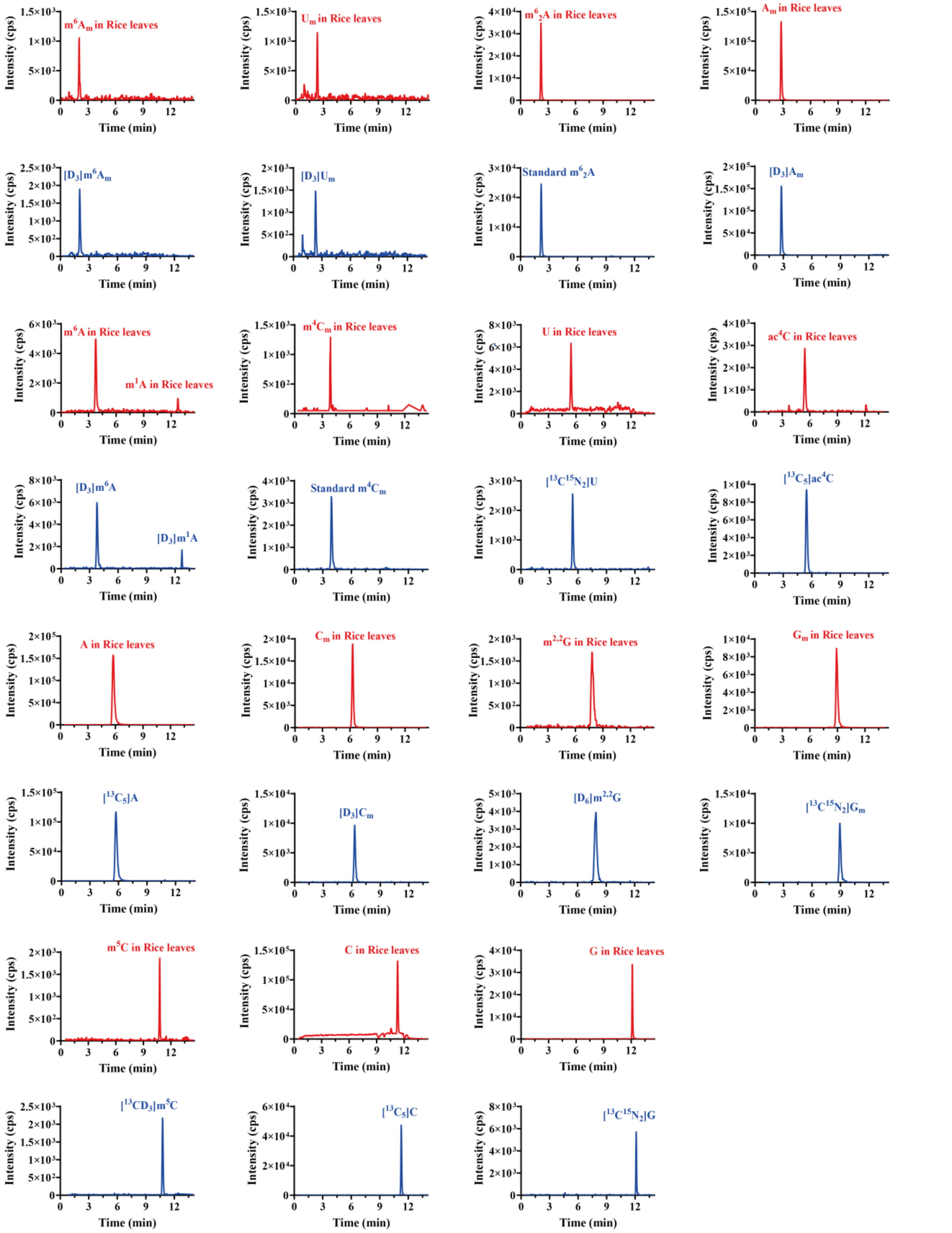


**Fig.S7** Identification of 16 kinds of nucleosides ($m^6A_m$, $U_m$, $m^6{}_2A$, $A_m$, $m^6A$, $m^1A$, $m^4C_m$, rU, $ac^4C$, rA, $C_m$, $m^{2,2}G$, $G_m$, $m^5C$, rC, and rG) in rice leaves samples by HILIC-MS/MS analysis. The extracted-ion chromatograms of the modifications detected in total RNA of rice leaves samples are highlighted in red and the spiked stable isotope-labeled internal standards are highlighted in blue.

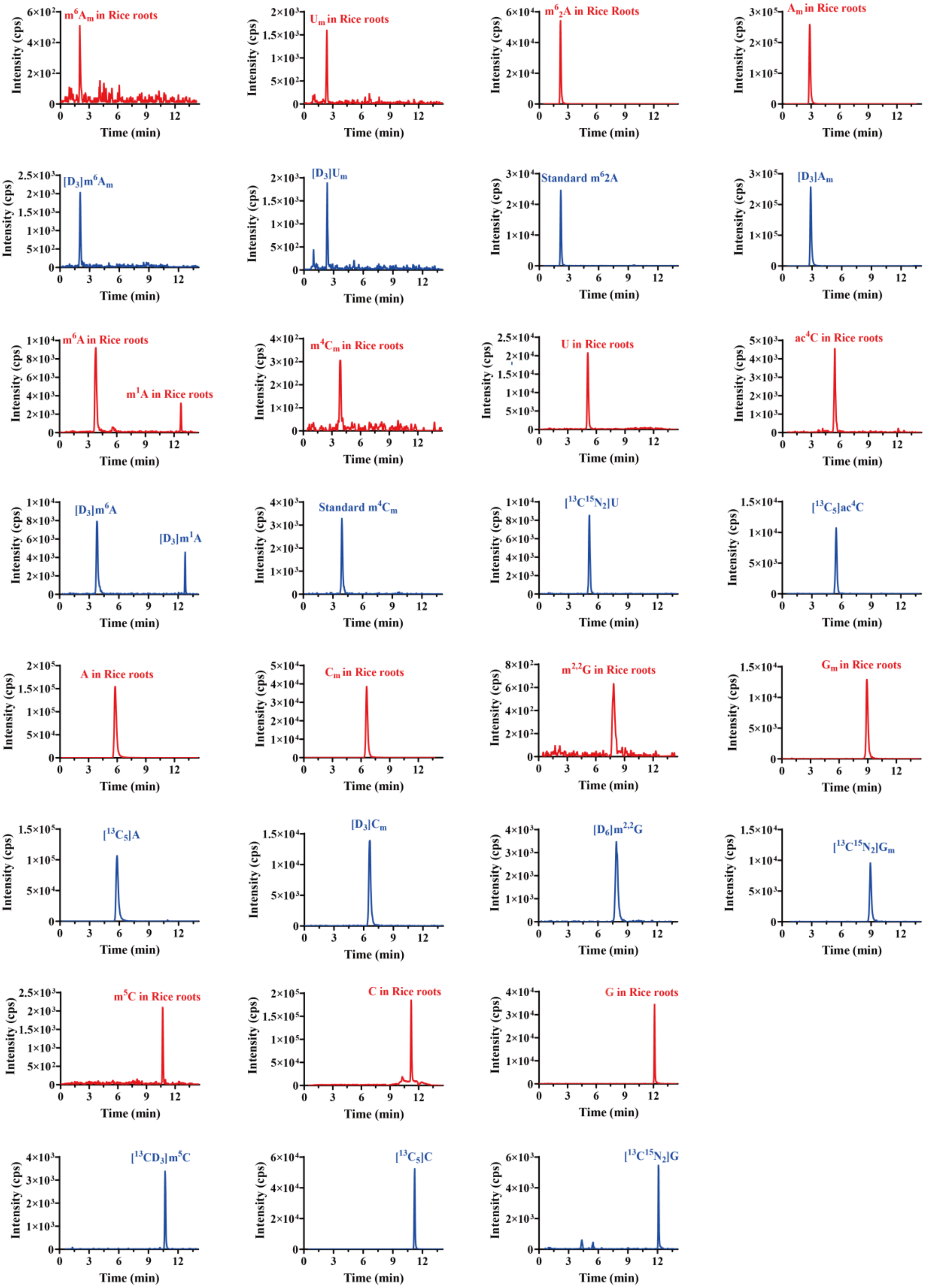


**Fig.S8** Identification of 16 kinds of nucleosides ($m^6A_m$, $U_m$, $m^6{}_2A$, $A_m$, $m^6A$, $m^1A$, $m^4C_m$, rU, $ac^4C$, rA, $C_m$, $m^{2,2}G$, $G_m$, $m^5C$, rC, and rG) in rice roots samples by HILIC-MS/MS analysis. The extracted-ion chromatograms of the modifications detected in total RNA of rice roots samples are highlighted in red and the spiked stable isotope-labeled internal standards are highlighted in blue.

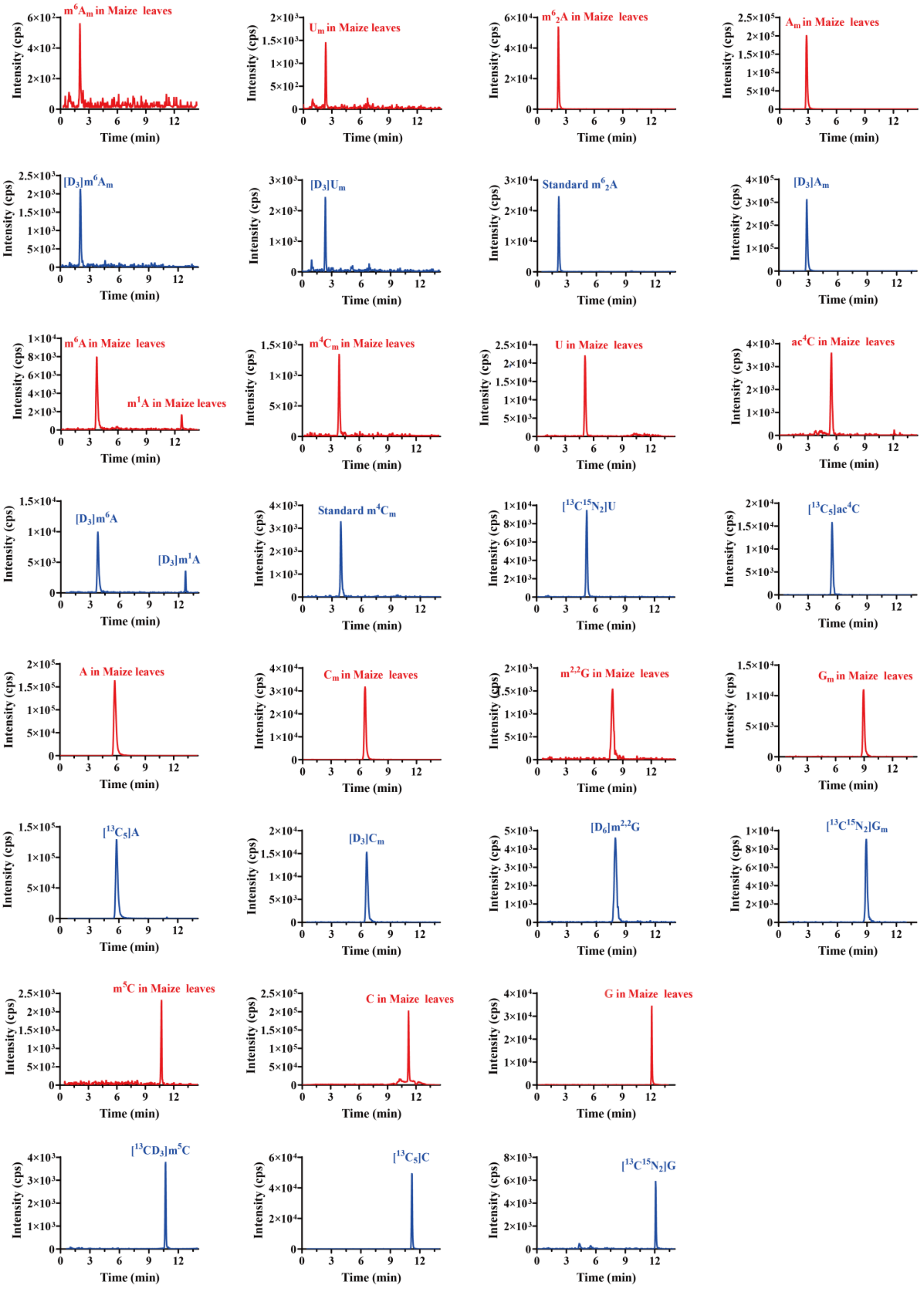


**Fig.S9** Identification of 16 kinds of nucleosides ($m^6A_m$, $U_m$, $m^6{}_2A$, $A_m$, $m^6A$, $m^1A$, $m^4C_m$, rU, $ac^4C$, rA, $C_m$, $m^{2,2}G$, $G_m$, $m^5C$, rC, and rG) in maize leaves samples by HILIC-MS/MS analysis. The extracted-ion chromatograms of the modifications detected in total RNA of maize leaves samples are highlighted in red and the spiked stable isotope-labeled internal standards are highlighted in blue.

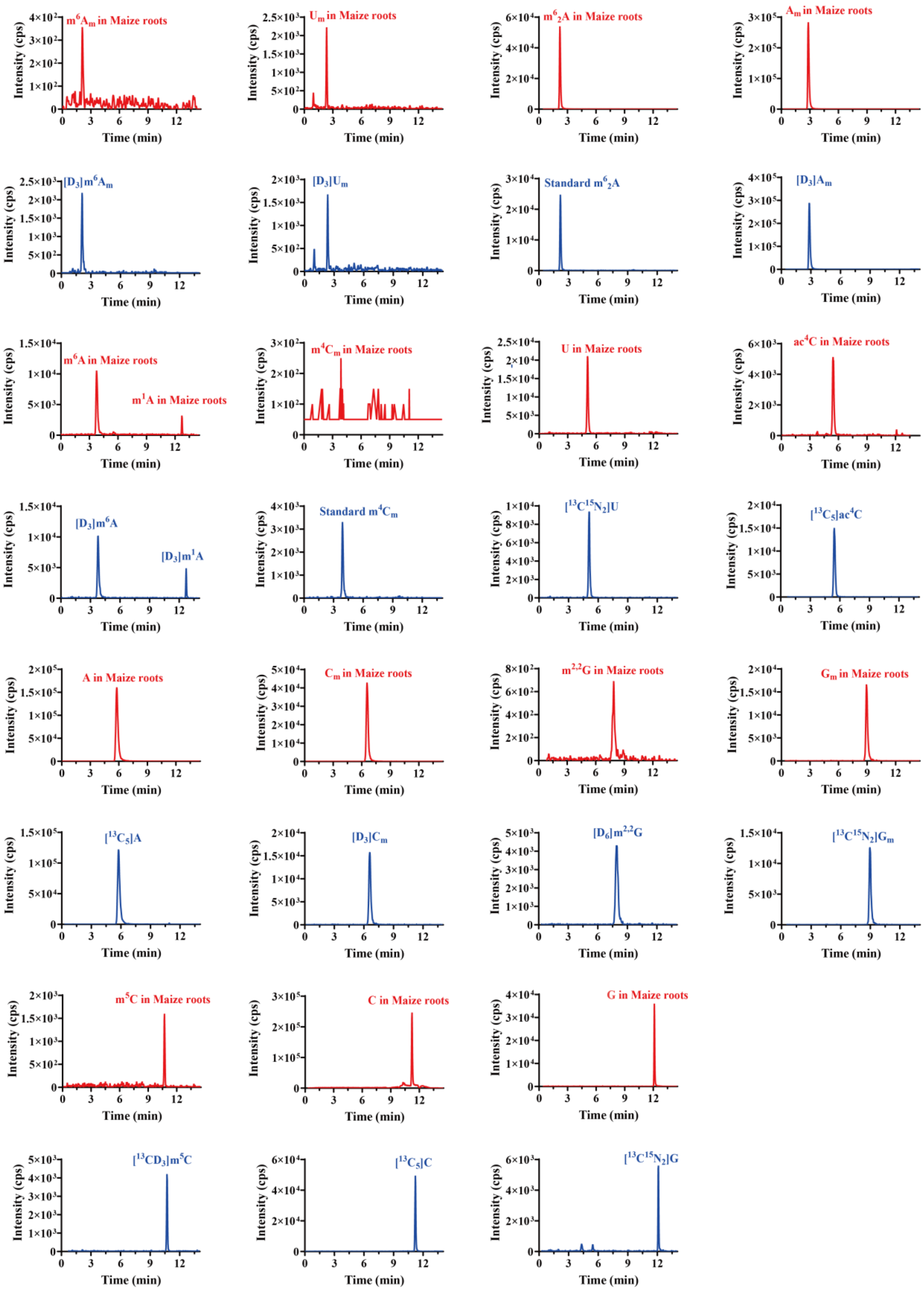


**Fig.S10** Identification of 16 kinds of nucleosides ($m^6A_m$, $U_m$, $m^6{}_2A$, $A_m$, $m^6A$, $m^1A$, $m^4C_m$, rU, $ac^4C$, rA, $C_m$, $m^{2,2}G$, $G_m$, $m^5C$, rC, and rG) in maize roots samples by HILIC-MS/MS analysis. The extracted-ion chromatograms of the modifications detected in total RNA of maize roots samples are highlighted in red and the spiked stable isotope-labeled internal standards are highlighted in blue.

**Table S1.** The information of 16 nucleoside standards and 14 isotopic nucleoside standards（For No. 1-17 obtained from Sigma Aldrich; 18-30 obtained from Toronto Research Chemical).

| No. | Nucleosides | Abbreviation | Molecular formula | Molecular weight |
|---|---|---|---|---|
| 1 | *N*6, 2′-O-dimethyladenosine | $m^6A_m$ | $C_{12}H_{17}N_5O_4$ | 295.29 |
| 2 | 2′-O-methyluridine | $U_m$ | $C_{10}H_{14}N_2O_6$ | 258.23 |
| 3 | *N*6, *N*6-dimethyladenosine | $m^6{}_2A$ | $C_{12}H_{17}N_5O_4$ | 295.29 |
| 4 | 2′-O-methyladenosine | $A_m$ | $C_{11}H_{15}N_5O_4$ | 281.27 |
| 5 | *N*6-methyladenosine | $m^6A$ | $C_{11}H_{15}N_5O_4$ | 281.27 |
| 6 | *N*1-methyladenosine | $m^1A$ | $C_{11}H_{15}N_5O_4$ | 281.27 |
| 7 | *N*4, 2′-O-dimethylcytidine | $m^4C_m$ | $C_{11}H_{17}N_3O_5$ | 271.27 |
| 8 | uridine | rU | $C_9H_{12}N_2O_6$ | 244.20 |
| 9 | *N*4-acetylcytidine | $ac^4C$ | $C_{11}H_{15}N_3O_6$ | 285.25 |
| 10 | adenosine | rA | $C_{10}H_{13}N_5O_4$ | 267.24 |
| 11 | 2′-O-methylcytidine | $C_m$ | $C_{10}H_{15}N_3O_5$ | 257.24 |
| 12 | *N*2, *N*2-dimethylguanosine | $m^{2,2}G$ | $C_{12}H_{17}N_5O_5$ | 311.29 |
| 13 | 2′-O-methylguanosine | $G_m$ | $C_{11}H_{15}N_5O_5$ | 297.27 |
| 14 | 5-methylcytidine | $m^5C$ | $C_{10}H_{15}N_3O_5$ | 257.24 |
| 15 | cytidine | rC | $C_9H_{13}N_3O_5$ | 243.22 |
| 16 | guanosine | rG | $C_{10}H_{13}N_5O_5$ | 283.24 |
| 17 | *N*6, 2′-O-dimethyladenosine-$D_3$ | $[D_3]m^6A_m$ | $C_{12}H_{14}D_3N_5O_4$ | 298.31 |
| 18 | 2′-O-methyluridine-$D_3$ | $[D_3]U_m$ | $C_{10}H_{11}D_3N_2O_6$ | 261.25 |
| 19 | 2′-O-methyladenosine-$D_3$ | $[D_3]A_m$ | $C_{11}H_{12}D_3N_5O_4$ | 281.27 |
| 20 | *N*6-methyladenosine-$D_3$ | $[D_3]m^6A$ | $C_{11}H_{12}D_3N_5O_4$ | 281.27 |
| 21 | *N*1-methyladenosine-$D_3$ | $[D_3]m^1A$ | $C_{11}H_{12}D_3N_5O_4$ | 281.27 |
| 22 | uridine-$^{13}C^{15}N_2$ | $[^{13}C^{15}N_2]rU$ | $C_8{}^{13}CH_{12}{}^{15}N_2O_6$ | 247.18 |
| 23 | *N*4-acetylcytidine-$^{13}C_5$ | $[^{13}C_5]ac^4C$ | $^{13}C_5$ $C_6$ $H_{15}$ $N_3$ $O_6$ | 290.28 |
| 24 | adenosine-$^{13}C_5$ | $[^{13}C_5]rA$ | $C_5{}^{13}C_5H_{13}N_5O_4$ | 272.20 |
| 25 | 2′-O-methylcytidine-$D_3$ | $[D_3]C_m$ | $C_{10}H_{12}D_3N_3O_5$ | 260.26 |
| 26 | *N*2, *N*2-dimethylguanosine-$D_6$ | $[D_6]m^{2,2}G$ | $C_{12}H_{11}D_6N_5O_5$ | 317.33 |
| 27 | 2′-O-methylguanosine-$D_3$ | $[D_3]G_m$ | $C_{11}H_{12}D_3N_5O_5$ | 300.29 |
| 28 | 5-methylcytidine-$^{13}CD_3$ | $[^{13}CD_3]m^5C$ | $C_9{}^{13}CH_{12}D_3N_3O_5$ | 261.25 |
| 29 | cytidine-$^{13}C_5$ | $[^{13}C_5]rC$ | $C_4{}^{13}C_5H_{13}N_3O_5$ | 248.18 |
| 30 | guanosine-$^{13}C^{15}N_2$ | $[^{13}C^{15}N_2]G$ | $C_9{}^{13}CH_{13}N_3{}^{15}N_2O_5$ | 286.22 |

**Table S2.** Identified modifications and relevant information in this study.

| Types | First Reported Date | First Reported method | First Reported Species | Reported in plant |
|---|---|---|---|---|
| **$m^6_2A$** | 1958[1] | Chromatographic and Electrophoretic | rat | 1982[2] |
| **$m^{2,2}G$** | 1958[3] | Paper chromatography and Electrophoresis | yeast | 1979[4] |
| **$m^5C$** | 1958[5] | Ultraviolet spectral | rat | 1982[2] |
| **$m^1A$** | 1961[6] | Chromatography and Ultraviolet spectral | yeast | 1979[4] |
| **$A_m$** | 1964[7] | Paper Chromatography | *Escherichia coli* | 1974[8] |
| **$G_m$** | 1964[7] | Paper Chromatography | *Escherichia coli* | 1974[8] |
| **$C_m$** | 1964[7] | Paper Chromatography | *Escherichia coli* | 1974[8] |
| **$U_m$** | 1964[7] | Paper Chromatography | *Escherichia coli* | 1974[8] |
| **$m^4C_m$** | 1966[9] | Ultraviolet spectral | *Escherichia coli* | in this work |
| **$ac^4C$** | 1966[10] | Chromatography and Electrophoresis | yeast | 1982[2] |
| **$m^6A$** | 1974[11] | High performance liquid chromatography | Novikoff hepatoma cells | 1979[12] |
| **$m^6A_m$** | 1975[13] | Thin-layer electrophoresis and Paper chromatography | HeLa cells | 2022[14] |

**Table S3**. Calibration curves, limits of detection (LODs) and limits of quantification (LOQs) for the analysis of RNA modifications.

| Compound | Linear Equation | $R^2$ Value | Linear Range (nM) | LOD (pM) | LOQ (pM) |
|---|---|---|---|---|---|
| **$m^6A_m$** | y = 2.918x - 0.0184 | 0.9994 | 0.025-1 | 10 | 25 |
| **$U_m$** | y = 0.0389x - 0.0039 | 0.9992 | 6.25-125 | 1250 | 2500 |
| **$m^6{}_2A$** | y = 0.829x - 0.0697 | 0.9998 | 1-50 | 5 | 25 |
| **$A_m$** | y = 1.1121x + 6.7255 | 0.9993 | 20-1000 | 25 | 50 |
| **$m^6A$** | y = 0.4693x - 0.0339 | 0.9998 | 0.500-20 | 25 | 50 |
| **$m^1A$** | y = 0.2893x + 0.0196 | 0.9998 | 0.500-25 | 1 | 5 |
| **$m^4C_m$** | y = 1.1663x - 0.0166 | 0.9993 | 0.050-5 | 10 | 25 |
| **rU** | y = 2.1912x + 0.1515 | 0.9995 | 250-10000 | 5000 | 10000 |
| **$ac^4C$** | y = 0.0242x + 0.0277 | 0.9996 | 2.5-100 | 25 | 50 |
| **rA** | y = 0.5464x + 0.0762 | 0.9998 | 500-25000 | 500 | 1000 |
| **$C_m$** | y = 0.1301x - 0.079 | 0.9994 | 5-250 | 2.5 | 5 |
| **$m^{2,2}G$** | y = 0.377x + 0.0377 | 0.9995 | 0.100-20 | 2.5 | 5 |
| **$G_m$** | y =0.0397x - 0.0883 | 0.9990 | 10-125 | 25 | 50 |
| **$m^5C$** | y = 0.0269x + 0.005 | 1 | 2.5-250 | 50 | 125 |
| **rC** | y = 2.0293x + 0.4623 | 0.9998 | 100-10000 | 100 | 500 |
| **rG** | y = 1.9308x + 0.3922 | 0.9994 | 500-20000 | 2500 | 500 |

**Table S4.** Accuracy and precision for the detection of RNA modifications.

| QC | Theoretical values (nM) | Intra-day (n=9) | | | Inter-day (n=3) | | |
|---|---|---|---|---|---|---|---|
| | | Mean ± SD （nM) | RSD (%) | Accuracy (%) | Mean ± SD （nM) | RSD (%) | Accuracy (%) |
| | 0.08 (Low) | 0.09 ± 0 | 2.22% | 109.96% | 0.09 ± 0 | 4.14% | 106.56% |
| $m^6A_m$ | 0.20 (Medium) | 0.22 ± 0 | 2.12% | 107.94% | 0.21 ± 0.01 | 3.65% | 104.71% |
| | 0.50 (High) | 0.54 ± 0.01 | 1.23% | 107.01% | 0.54 ± 0.01 | 2.33% | 107.59% |
| | 12 (Low) | 12.03 ± 0.26 | 2.17% | 100.25% | 11.95 ± 0.62 | 5.21% | 99.54% |
| $U_m$ | 30 (Medium) | 27.67 ± 0.71 | 2.56% | 92.22% | 27.43 ± 0.15 | 0.56% | 91.42% |
| | 75 (High) | 70.76 ± 0.85 | 1.20% | 94.35% | 71.49 ± 2.65 | 3.70% | 95.32% |
| | 2.5 (Low) | 2.32 ± 0.08 | 3.54% | 92.69% | 2.32 ± 0.04 | 1.93% | 92.83% |
| $m^6_2A$ | 6.25 (Medium) | 5.59 ± 0.16 | 2.83% | 111.77% | 5.81 ± 0.1 | 1.68% | 92.98% |
| | 15.625 (High) | 14.96 ± 0.23 | 1.56% | 95.73% | 15.12 ± 0.2 | 1.33% | 96.79% |
| | 75 (Low) | 75.48 ± 0.98 | 1.29% | 100.64% | 74.11 ± 0.46 | 0.62% | 98.81% |
| $A_m$ | 120 (Medium) | 123.41 ± 0.55 | 0.45% | 102.84% | 122.66 ± 1.07 | 0.87% | 102.21% |
| | 300 (High) | 293.33 ± 2.26 | 0.77% | 97.78% | 296.5 ± 3.11 | 1.05% | 98.83% |
| | 1.25 (Low) | 1.28 ± 0.02 | 1.61% | 102.66% | 1.26 ± 0.04 | 3.40% | 100.47% |
| $m^6A$ | 2 (Medium) | 2.14 ± 0.15 | 6.83% | 106.88% | 1.95 ± 0.05 | 2.73% | 97.63% |
| | 5 (High) | 5.17 ± 0.05 | 1.05% | 103.33% | 5.2 ± 0.2 | 3.91% | 103.94% |
| | 1.25 (Low) | 1.28 ± 0.07 | 5.66% | 102.19% | 1.38 ± 0.11 | 7.77% | 110.15% |
| $m^1A$ | 2 (Medium) | 1.89 ± 0.11 | 5.72% | 94.49% | 1.91 ± 0.02 | 0.81% | 95.72% |
| | 5 (High) | 4.37 ± 0.12 | 2.83% | 87.45% | 4.51 ± 0.16 | 3.56% | 90.20% |
| | 0.2 (Low) | 0.21 ± 0.01 | 3.56% | 103.73% | 0.21 ± 0 | 1.83% | 103.55% |
| $m^4C_m$ | 0.5 (Medium) | 0.49 ± 0.03 | 5.54% | 97.35% | 0.48 ± 0.01 | 2.27% | 96.20% |
| | 1.25 (High) | 1.22 ± 0.06 | 4.67% | 97.58% | 1.23 ± 0.03 | 2.77% | 98.51% |
| | 1000 (Low) | 996.05 ± 26.83 | 2.69% | 99.60% | 961.79 ± 7.76 | 0.81% | 96.18% |
| rU | 2500 (Medium) | 2390.71 ± 71.12 | 2.97% | 95.63% | 2478.16 ± 69.28 | 2.80% | 99.13% |
| | 6250 (High) | 6299.27 ± 237.93 | 3.78% | 100.79% | 6205.08 ± 223.34 | 3.60% | 99.28% |
| | 0.8 (Low) | 0.75 ± 0.01 | 1.11% | 93.73% | 0.76 ± 0.02 | 2.22% | 94.90% |
| $ac^4C$ | 2 (Medium) | 1.92 ± 0.02 | 1.21% | 96.10% | 1.94 ± 0.02 | 0.79% | 97.12% |
| | 5 (High) | 5.19 ± 0.03 | 0.53% | 103.82% | 5.2 ± 0.01 | 0.23% | 103.98% |
| | 1600 (Low) | 1535.95 ± 3.52 | 0.23% | 96.00% | 1537.18 ± 17.28 | 1.12% | 96.07% |
| rA | 4000 (Medium) | 3940.18 ± 21.11 | 0.54% | 98.50% | 3969.56 ± 73.64 | 1.86% | 99.24% |
| | 10000 (High) | 10116.39 ± 21.11 | 0.21% | 101.16% | 10145.64 ± 119.37 | 1.18% | 101.46% |
| | 12 (Low) | 12.46 ± 0.07 | 0.58% | 103.80% | 12.43 ± 0.06 | 0.44% | 103.55% |
| $C_m$ | 30 (Medium) | 28.39 ± 0.12 | 0.42% | 94.62% | 28.43 ± 0.33 | 1.18% | 94.78% |
| | 75 (High) | 73.07 ± 0.95 | 1.31% | 97.43% | 72.71 ± 0.98 | 1.35% | 96.95% |
| | 1.6 (Low) | 1.47 ± 0.01 | 0.86% | 92.05% | 1.49 ± 0.04 | 2.96% | 92.91% |
| $m^{2,2}G$ | 4 (Medium) | 3.55 ± 0.03 | 0.84% | 88.69% | 3.59 ± 0.07 | 1.93% | 89.68% |
| | 10 (High) | 9.62 ± 0.07 | 0.76% | 96.18% | 9.64 ± 0.2 | 2.06% | 96.42% |
| | 16 (Low) | 17.7 ± 0.36 | 2.05% | 110.64% | 17.19 ± 0.23 | 1.36% | 107.44% |
| $G_m$ | 40 (Medium) | 40.37 ± 0.19 | 0.46% | 100.92% | 40.53 ± 0.44 | 1.10% | 101.32% |
| | 100 (High) | 100.9 ± 1.57 | 1.56% | 100.90% | 101.41 ± 0.85 | 0.84% | 101.41% |

**Table S4 Continued.**

| QC | Theoretical values (nM) | Intra-day (n=9) | | | Inter-day (n=3) | | |
|---|---|---|---|---|---|---|---|
| | | Mean ± SD （nM) | RSD (%) | Accuracy (%) | Mean ± SD （nM) | RSD (%) | Accuracy (%) |
| $m^5C$ | 2 (Low) | 1.88 ± 0.02 | 0.86% | 94.06% | 1.9 ± 0.05 | 2.72% | 95.06% |
| | 5 (Medium) | 4.57 ± 0.08 | 1.67% | 91.32% | 4.61 ± 0.05 | 1.19% | 92.25% |
| | 12.5 (High) | 11.75 ± 0.07 | 0.59% | 94.00% | 11.68 ± 0.15 | 1.26% | 93.45% |
| rC | 2000 (Low) | 1977.7 ± 26.76 | 1.35% | 98.88% | 2015.51 ± 39.56 | 1.96% | 100.78% |
| | 5000 (Medium) | 4638.34 ± 42.77 | 0.92% | 92.77% | 4650.23 ± 85.6 | 1.84% | 93.00% |
| | 12000 (High) | 11053.83 ± 90.84 | 0.82% | 92.12% | 11121.46 ± 115.91 | 1.04% | 92.68% |
| rG | 1600 (Low) | 1605.93 ± 17.55 | 1.09% | 100.37% | 1594.25 ± 19 | 1.19% | 99.64% |
| | 4000 (Medium) | 4068.98 ± 41.07 | 1.01% | 101.72% | 4074.5 ± 135.49 | 3.33% | 101.86% |
| | 10000 (High) | 10131.44 ± 125.51 | 1.24% | 101.31% | 10209.51 ± 231.52 | 2.27% | 103.83% |

**Table S5.** Measured average levels of RNA modifications in total RNA upon NAA treatment from *Marchantia polymorpha* and *Arabidopsis thaliana*.

| % | ***Marchantia Polymorpha*** | | ***Arabidopsis Thaliana*** | |
|---|---|---|---|---|
| | **Control** | **NAA Treatment** | **Control** | **NAA Treatment** |
| **$m^6A_m$/A** | ND | ND | 0.0026 ± 0.0001 | 0.0024 ± 0.0002 |
| **$U_m$/U** | 2.0603 ± 0.1650 | 2.2281 ± 0.1011 | 2.2250 ± 0.1164 | 2.7562 ± 0.2610 |
| **$m^6_2A$/A** | 0.2144 ± 0.0068 | 0.2068 ± 0.0052 | 0.1946 ± 0.0061 | 0.2050 ± 0.0031 |
| **$A_m$/A** | 4.6542 ± 0.0811 | 4.6400 ± 0.0852 | 4.2718 ± 0.1016 | 4.3294 ± 0.146 |
| **$m^6A$/A** | 0.0798 ± 0.0024 | 0.0775 ± 0.0011 | 0.0843 ± 0.0027 | 0.0867 ± 0.0017 |
| **$m^1A$/A** | 0.0748 ± 0.0021 | 0.0761 ± 0.0053 | 0.0892 ±0.0099 | 0.0823 ± 0.00608 |
| **$m^4C_m$/C** | 0.014 ± 0.0022 | 0.0136 ± 0.0016 | 0.0178 ± 0.0015 | 0.0125 ± 0.0019 |
| **$ac^4C$/C** | 0.0729 ± 0.0083 | 0.0472 ± 0.0041 | 0.0737 ± 0.0017 | 0.0656 ± 0.0032 |
| **$C_m$/C** | 1.4190 ± 0.1095 | 1.1881 ± 0.1793 | 1.0285 ± 0.1975 | 0.9876 ± 0.1981 |
| **$m^{2,2}G$/G** | 0.0210 ± 0.0014 | 0.0200 ± 0.0008 | 0.0231 ± 0.0019 | 0.0193 ± 0.0006 |
| **$G_m$/G** | 1.2134 ± 0.0341 | 1.1761 ± 0.0398 | 1.2777 ± 0.0475 | 1.3560 ± 0.0240 |
| **$m^5C$/C** | 0.1161 ± 0.0083 | 0.1026 ± 0.0147 | 0.1163 ± 0.0047 | 0.0940 ± 0.0071 |

Data present the mean ± standard error of the mean (SEM) of three independent experiments. ND, not detected.

**Table S6.** Measured average levels of RNA modifications in total RNA upon NAA treatment from rice and maize.

| % | Rice Leaves | | Rice Roots | | Maize Leaves | | Maize Roots | |
|---|---|---|---|---|---|---|---|---|
| | Control | NAA Treatment | Control | NAA Treatment | Control | NAA Treatment | Control | NAA Treatment |
| $m^6A_m/A$ | 0.0057 ± 0.0006 | 0.0028 ± 0.0002 | 0.0036 ± 0.0002 | 0.0039 ± 0.0004 | 0.0031 ± 0.0002 | 0.0027 ± 0.0002 | <LOQ | <LOQ |
| $U_m/U$ | 1.7718 ± 0.1391 | 1.7654 ± 0.1665 | 2.5797 ± 0.1115 | 2.7512 ± 0.1722 | 1.9560 ± 0.2046 | 2.4157 ± 0.1765 | 2.9484 ± 0.1287 | 2.5185 ± 0.2634 |
| $m^6{}_2A/A$ | 0.1844 ± 0.0085 | 0.1763 ± 0.0042 | 0.2127 ± 0.0046 | 0.2108 ± 0.0045 | 0.1782 ± 0.0039 | 0.1923 ± 0.0055 | 0.2063 ± 0.0059 | 0.2243 ± 0.006 |
| $A_m/A$ | 3.5857 ± 0.0943 | 3.4302 ± 0.0509 | 4.9987 ± 0.0645 | 5.0979 ± 0.1253 | 3.3054 ± 0.0786 | 4.0107 ± 0.0573 | 5.1861 ± 0.0985 | 5.5853 ± 0.1464 |
| $m^6A/A$ | 0.0726 ± 0.0027 | 0.0639 ± 0.0011 | 0.1058 ± 0.0012 | 0.1061 ± 0.0019 | 0.0702 ± 0.0014 | 0.0813 ± 0.0021 | 0.0988 ± 0.0036 | 0.1125 ± 0.0019 |
| $m^1A/A$ | 0.08152 ± 0.0053 | 0.0666 ± 0.0055 | 0.1278 ± 0.0065 | 0.1035 ±0.0059 | 0.0793 ±0.0054 | 0.1053 ±0.0192 | 0.1012 ±0.0046 | 0.1531 ±0.0149 |
| $m^4C_m/C$ | 0.0267 ± 0.0021 | 0.0276 ± 0.0026 | 0.0069 ± 0.0005 | 0.0077 ± 0.0004 | 0.0185 ± 0.0017 | 0.0240 ± 0.0051 | ND | ND |
| $ac^4C/C$ | 0.0532 ± 0.0013 | 0.0511 ± 0.0028 | 0.0723 ± 0.0061 | 0.0719 ± 0.007 | 0.0550 ± 0.0056 | 0.0629 ± 0.0023 | 0.0693 ± 0.0078 | 0.0727 ± 0.0048 |
| $C_m/C$ | 0.9679 ± 0.0582 | 0.8966 ± 0.0617 | 1.2906 ± 0.0419 | 1.2706 ± 0.0437 | 0.8540 ± 0.0414 | 1.1414 ± 0.0410 | 1.2235 ± 0.1391 | 1.2347 ± 0.0337 |
| $m^{2,2}G/G$ | 0.0340 ± 0.0029 | 0.0314 ± 0.0009 | 0.0139 ± 0.0008 | 0.0148 ± 0.0020 | 0.0348 ± 0.0010 | 0.0326 ± 0.0027 | 0.0083 ± 0.0006 | 0.0171 ± 0.0012 |
| $G_m/G$ | 0.9439 ± 0.0462 | 0.9925 ± 0.038 | 1.2317 ± 0.0525 | 1.2947 ± 0.0546 | 0.9804 ± 0.0516 | 1.0926 ± 0.0351 | 1.2839 ± 0.0419 | 1.3146 ± 0.0359 |
| $m^5C/C$ | 0.1460 ± 0.0086 | 0.1109 ± 0.0077 | 0.1242 ± 0.0079 | 0.1124 ± 0.0052 | 0.1066 ± 0.0056 | 0.1259 ± 0.0064 | 0.0902 ± 0.0125 | 0.0991 ± 0.0059 |

Data present the mean ± standard error of the mean (SEM) of three independent experiments.<LOQ: below the instrumental limit of detection. ND, not detect.

**Table S7.** The MRM transitions and optimal parameters for the analysis of nucleosides by mass spectrometry.

| NO. | Compound | MRM ion transition (m/z) | CE (V) | DP (V) | EP (V) | CXP (V) |
|---|---|---|---|---|---|---|
| **1** | $m^6A_m$ | 296.106→150 | 25 | 50 | 10 | 10 |
| **2** | $U_m$ | 259.1→112 | 18 | 60 | 6 | 10 |
| **3** | $m^6{}_2A$ | 296.1→164 | 28 | 55 | 10 | 16 |
| **4** | $A_m$ | 282.1→136 | 22 | 50 | 9 | 16 |
| **5** | $m^6A$ | 282.106→150 | 26 | 50 | 12 | 12 |
| **6** | $m^1A$ | 282.101→150 | 23 | 50 | 6 | 10 |
| **7** | $m^4C_m$ | 272.104→126 | 19 | 50 | 7 | 10 |
| **8** | rU | 245.1→113 | 1 5 | 60 | 10 | 10 |
| **9** | $ac^4C$ | 286.1→154 | 16 | 45 | 10 | 10 |
| **10** | rA | 268.1→136 | 10 | 50 | 10 | 10 |
| **11** | $C_m$ | 258.1→112 | 16 | 45 | 10 | 9 |
| **12** | $m^{2,2}G$ | 312.1→180 | 20 | 50 | 6 | 5 |
| **13** | $G_m$ | 298.1→152 | 16 | 50 | 7 | 5 |
| **14** | $m^5C$ | 258.105→126 | 16 | 50 | 6 | 10 |
| **15** | C | 244.1→112 | 16 | 45 | 10 | 10 |
| **16** | G | 284.1→152 | 20 | 50 | 10 | 10 |
| **17** | $[D_3]m^6A_m$ | 299.106→153 | 14 | 50 | 4 | 10 |
| **18** | $[D_3]U_m$ | 262.1→113 | 18 | 60 | 6 | 8 |
| **19** | $[D_3]A_m$ | 285.1→136 | 22 | 50 | 9 | 8 |
| **20** | $[D_3]m^6A$ | 285.106→153 | 26 | 50 | 12 | 14 |
| **21** | $[D_3]m^1A$ | 285.101→153 | 23 | 50 | 6 | 10 |
| **22** | $[^{13}C^{15}N_2]rU$ | 248.1→116 | 15 | 60 | 10 | 14 |
| **23** | $[^{13}C_5]ac4C$ | 291.1→154 | 16 | 45 | 10 | 12 |
| **24** | $[^{13}C_5]rA$ | 273.1→136 | 23 | 50 | 10 | 12 |
| **25** | $[D_3]C_m$ | 261.1→112 | 16 | 45 | 10 | 8 |
| **26** | $[D_6]m^{2,2}G$ | 318.1→186 | 20 | 50 | 6 | 8 |
| **27** | $[D_3]G_m$ | 301.1→152 | 16 | 50 | 7 | 7 |
| **28** | $[^{13}CD_3]m^5C$ | 262.105→130 | 16 | 50 | 6 | 7 |
| **29** | $[^{13}C_5]rC$ | 249.1→112 | 20 | 45 | 10 | 10 |
| **30** | $[^{13}C^{15}N_2]rG$ | 287.1→155 | 20 | 50 | 10 | 10 |

*Commun.* **2022,** *13* (1), 5636.
